# Dermal fibroblast cultures recapitulate differences between deermice and mice in responses to a Toll-like receptor agonist

**DOI:** 10.1101/2025.07.16.665222

**Authors:** Jonathan V. Duong, Aqsa Motiwala, William J. Hotz, Landen Gozashti, Anthony D. Long, Alan G. Barbour

## Abstract

The white-footed deermouse *Peromyscus leucopus* is a primary reservoir for the agents of Lyme disease and other zoonoses in North America and manifests infection tolerance for the bacteria, protozoa, and viruses it hosts. In previous in vivo studies *P. leucopus* and *M. musculus* differed in the degree of sickness and profiles of biomarkers after exposure to bacterial lipopolysaccharide, a TLR4 agonist. As an approach for assessing immunity of mammals in nature and for longitudinal studies of colony animals in the laboratory, we evaluated using bulk and single cell RNA-seq primary dermal fibroblast cultures of *P. leucopus* and *M. musculus* in their short-term responses to a TLR2 agonist lipopeptide. By single cell RNA-seq cultures of both species comprised at least two types of fibroblasts, which were further differentiated in their responses to TLR agonists. With continued passage the mouse cell population lost viability, while the deermouse cell population spontaneously transformed into a cell line stably maintained under standard conditions. Bulk RNA-seq revealed distinctive profiles for deermouse and mouse cells in arginine metabolism gene expression, high baseline transcription of the antioxidant transcription factor Nfe2l2 (Nrf2) in deermouse fibroblasts, and the transcription of the aging-associated cytokine interleukin-11 in agonist-treated mouse fibroblasts but not deermouse fibroblasts. In both species’ cultures there was increased transcription of several types of endogenous retrovirus (ERV) and transposable elements (TE) after exposure to the agonist. The transcribed ERV/TE sequences in *M. musculus* cells were generally longer in length and with greater potential for translation than sequences in treated *P. leucopus* cells. The results indicate feasibility of this in vitro model for both laboratory- and field-based studies and that inherent differences between deermice and mice in cell-autonomous innate immune responses and ERV/TE activation can be demonstrated in dermal fibroblasts as well as the animals themselves.

## Introduction

As Hagai et al. noted, the immune system’s characteristics of both rapid divergence and high cell-to-cell variability “seem to be at odds with strong regulatory constraints imposed on the host immune response: the need to execute a well-coordinated and carefully balanced program to avoid tissue damage and pathological immune conditions” (1). To resolve this apparent paradox, established animal models, principally the house mouse *Mus musculus*, are commonly enlisted. But what if the question could be more profitably addressed using another species (2), such as one that in its natural environment thrives while persistently infected with microbes of trans-species transmission potential? Ideally, wild populations of the species would be accessible for field-based studies, and there would be vivarium-bred stock colonies available for laboratory-based studies.

Our nominee as an alternative to the house mouse for this question is the North American species *Peromyscus leucopus*, the white-footed deermouse (3, 4). Mouse-like in appearance and size, this and other deermice, such as *P. maniculatus*, belong to the family Cricetidae, along with hamsters, voles, and woodrats, and not the family Muridae, the taxonomic clade of laboratory mice and rats (5). *P. leucopus* is abundant in eastern and central United States and found in a variety of environments. Closed colonies of genetically diverse stock exist at different institutions. As a foundation for forward and reverse genetics, as well as for bulk and single cell RNA-seq, there are high quality, chromosome-scale genome assemblies of both *P. leucopus* and *P. maniculatus*, with full annotation and millions of single nucleotide polymorphisms and other variants that have been catalogued to date (6, 7).

Another rationale for choosing *P. leucopus* is its public health importance as a reservoir for several agents of human zoonoses. Besides the Lyme disease agent, *P. leucopus* is also a natural host for the obligate intracellular bacterium of anaplasmosis, the apicomplexan protozoan of babesiosis, and the flavivirus of Powassan viral encephalitis. Deermice may harbor a given pathogen at loads sufficient for transmission to a vector, but they are largely free of disability or discernible effects on fitness (reviewed in (8)). Infected with the spirochete *Borreliella burgdorferi*, the principal agent of Lyme disease in North America, *P. leucopus* display few of the pathologic changes or inflammation observed in infected *M. musculus*.

This phenomenon is known as infection tolerance (9, 10), which is characterized by immunological and physiological adaptations that minimize the harm from a pathogen’s presence without necessarily reducing its burden. Infection tolerance by this definition subtly differs from “disease tolerance”, which can be thought of as the ability of a host to survive or maintain fitness despite experiencing disease symptoms or pathology (11-13). Disease tolerance is a concept that places more emphasis on damage mitigation and repair mechanisms. Whether we interpret a finding as tolerance of infection or of disease, the phenomenon may have relevance for studies of aging. *P. leucopus* has a maximum longevity that is two-to-three times longer than that of *M. musculus* (14, 15). Infection tolerance and greater longevity for their body sizes are characteristics *P. leucopus* has in common with some bats (16, 17).

These distinctions between deermice and mice inspired our comparative studies of these representatives of two rodent genera. The experimental study design was to induce a host response to a microbial product (18), specifically bacterial lipopolysaccharide (LPS), which is a Toll-like receptor 4 (TLR4) agonist and known to elicit acute systemic inflammation (19, 20). Thus treated *P. leucopus* had profiles of genome-wide RNA-seq of the blood, spleen, and liver that distinguished it from treated *M. musculu*s. In general, the LPS-treated deermice displayed alternatively-activated macrophage polarization rather than the expected classically-activated, mitigation of the effects of neutrophil activation, and restrained type 1 and type 2 interferon responses. More specifically, *P. leucopus* was distinguished from *M. musculus* by an inverted ratio of nitric oxide synthase 2 (Nos2) to arginase 1 (Arg1) transcription, remarkably high transcription of the Slpi gene for secretory leukocyte peptidase inhibitor (Slpi), and relatively diminished interferon expression.

For the present study of *P. leucopus* and *M. musculus*, we looked to reduce the subject of the experiment to one that retains characteristics of live animals, yet would also serve for studies of wild animals that could be captured and then released (21). Blood sampling from trapped then released animals is an established procedure for the study of the immunology or physiology of animals in nature (18). However, assays of whole blood or serum are limited to what a single volume of blood can provide and nothing more. If the spleen is the tissue (22), the capture is a terminal event for the animal. For field and laboratory work on Lyme disease a specimen commonly obtained has been punch biopsies of ear skin tissue, which are then subjected to culture or quantitative PCR for *B. burgdorferi* bacteria (23, 24).

It occurred to us that this biopsied tissue could also be propagated ex vivo in the laboratory, thus allowing for a much expanded set of experiments with further rounds of multiplication. A critique would hold that a culture of cells of a tissue is not evaluable as to its “fitness”, as one could for the organisms themselves in an ecology or evolutionary biology context. From this perspective, a tissue culture cannot manifest infection tolerance per se. But because infection tolerance is characterizable by specific adaptations (25), some of which are of the measurable sort with isolated tissues or cells, there is justification for proposing primary cultures as in vitro correlates of a phenomenon that applies primarily at the organismal or population level.

As a proof-of-principle for *Peromyscus*, we cultivated primary dermal fibroblasts obtained from the ears of heterogenous stock *P. leucopus* or outbred *M. musculus* in a laboratory setting. Low-passage cultures of skin cells were then exposed to either TLR2 agonist or buffer alone. TLR2, as a heterodimer with TLR1, is the pattern recognition receptor (PRR) for pathogen-associated molecular pattern (PAMP) represented by the bacterial lipoproteins of *B. burgdorferi* (26). The resultant specimens were subjected to bulk and single cell RNA-seq, for which the reference sets were genome-wide protein coding sequences of *P. leucopus* and *M. musculus*, as well as full sets of endogenous retrovirus-derived elements of each species (27). We also characterized a spontaneously transformed line of *P. leucopus* dermal fibroblasts that continued in its capacity for proliferation long after an analogous *M. musculus* fibroblast line had ceased to replicate.

## Results

### Primary cultures and serial passages

For these experiments full-thickness samples of freshly-excised ears of euthanized *P. leucopus* or *M. musculus* animals were cultivated in a type of medium and under conditions long-used for primary fibroblast culture (28). There was no attempt to preserve the spatial characteristics of skin tissue by addition of growth factors or other supplements. The oxygen concentration was that of an incubator with 5% CO_2_ and at sea level without any adjustments to limit oxidative stress (29). Cells of the original cultures and then of subsequent passages were aliquoted and frozen, thereby preserving the history of the two lineages. Examples of early passage cells from *P. leucopus* and *M. musculus* in culture are shown in Figure 1. Successful cultivation of *P. leucopus* dermal fibroblasts was also achieved with single 2 mm punch biopsies of ears of animals.

**Figure 1.**
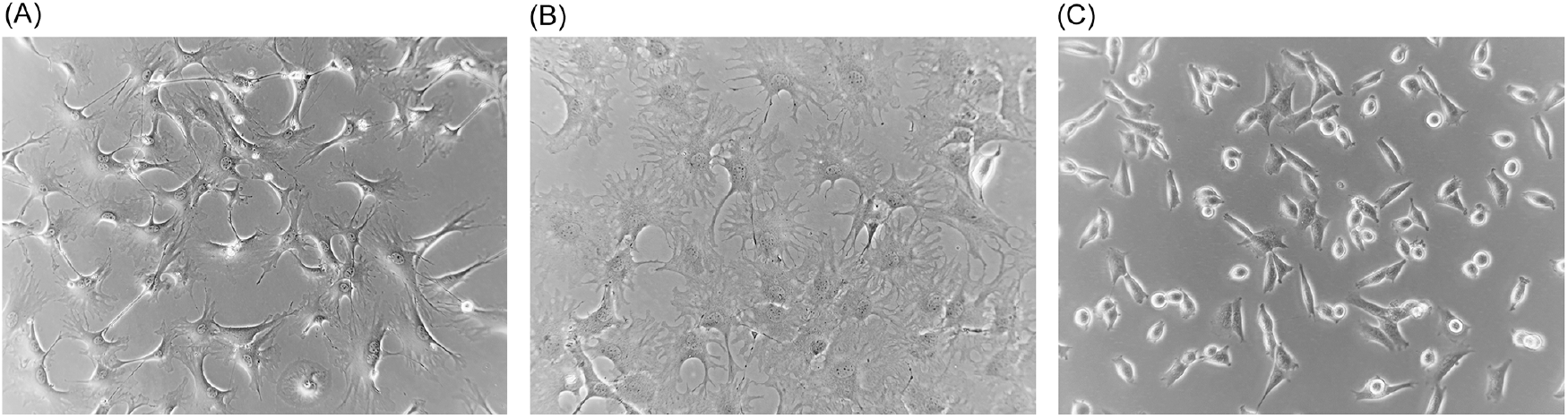
Phase microscopy photographs of cultures of dermal fibroblasts of *Mus musculus* (panel A) and *Peromyscus leucopus* (panels B and C) and. Panels (A) and (B) are of second passage cells of each species, and panel C is of high passage (P47) transformed cells. Magnification, 150x.

In their initial cultures *M. musculus* cells were confluent on the bottom of the dishes by day 5 to 7, while the *P. leucopus* cells at the same starting inoculum achieved that state only by day 10 to 12. Doubling times for early passage populations of *P. leucopus* cells and *M. musculus* cells were ∼56 h and ∼28 h, respectively. By passage 16 (or ∼70 doublings), the time for the *M. musculus* cultures to reach confluency had lengthened to 10-14 d, with a doubling time the same as the *P. leucopus* population at that point. Within a few more passages the *M. musculus* cells failed to adhere and growth further slowed. The emergence of an established cell line, as can occur with cultivation of mouse embryo cells (30), was not observed.

Under the same cultivation conditions, by passage 20 (∼86 doublings) the time to confluency for *P. leucopus* cells to reach confluency was only 6-8 days, which corresponded to a doubling time of 14 h. Thereafter the *P. leucopus* cells grew at this higher rate through more than 47 serial passages with no signs of abnormalities in adherence and growth noted for the mouse cells. This apparent adaptation of the *P. leucopus* cells to in vitro life was accompanied by a change in cell morphology; they became shorter and more rounded with fewer extensions while retaining their adherence capacity (Figure 1). The *P. leucopus* dermal fibroblast culture was considered “spontaneously transformed” thereafter (31).

### Bulk RNA-seq

For the experiments with the lipopeptide TLR agonist Pam3CSK4, the sources of tissue for in vitro cultivation were 5 adult outbred LL stock *P. leucopus* (3 females and 2 males) and 5 adult outbred CD-1 *M. musculus* (3 females and 2 males) (Table S1). Second passage (P2) cultures (∼16 doublings) at 80-90% confluency were split in three and then after growth for 24 h subjected to the following exposures for 4 h: no treatment control or the lipopeptide TLR2 agonist Pam3CSK4 at 1 µg/ml or 10µg/ml. The cDNA libraries were mRNA stranded and yielded ranges of 1.03-1.42 × 10^8^ paired-end 150 nt (PE150) reads for the 15 *P. leucopus* samples and 1.25-1.63 × 10^8^ reads for *M. musculus*. Reference sets were the protein coding sequences (CDS) of the genome of a female *P. leucopus* of LL stock and the reference C57BL/6 genome of a female *M. musculus*. As a consequence of the more comprehensive isoform annotation for the mouse genome to that point, there were four times as many CDS sequences listed for *M. musculus* as for *P. leucopus*. Accordingly, for comparability, the first listed isoform for *M. musculus* was used for the reference set, resulting in nonredundant CDS sets of 22,760 for *M. musculus* and 22,654 for *P. leucopus* (Dryad Tables D1 and D2).

Distributions of log-transformed TPM values by cumulative count of genes were similar between species (Figure S1). There were 14,979 genes in common to both species and that met the criterion of TPM ≥10 in at least one of the individual 30 samples (Dryad Table D3). There was also similarity between species in the log-log plots of *p* values and fold changes of paired samples; for both species the numbers of DEGs with higher transcription after agonist exposure were greater than the number of DEGs with reduced transcription post-exposure (Figure 2 panels A and B). The mean coefficients of determination (*R*^*2*^) values in log-transformed reads by CDS between paired values for the 1 µg and 10 µg per ml concentrations were 0.994 (0.992-0.996) for *M. musculus* and 0.964 (0.927-1.00) for *P. leucopus* (*p* = 0.15), indicating low to negligible dose effect within the 10-fold range of the study (Dryad Tables D1 and D2). Unless otherwise noted, comparisons with paired controls were done with cells exposed to the 1 µg/ml concentration, as likely more representative of the tissue environment in the *B. burgdorferi*-infected skin of *P. leucopus* reservoir hosts.

**Figure 2.**
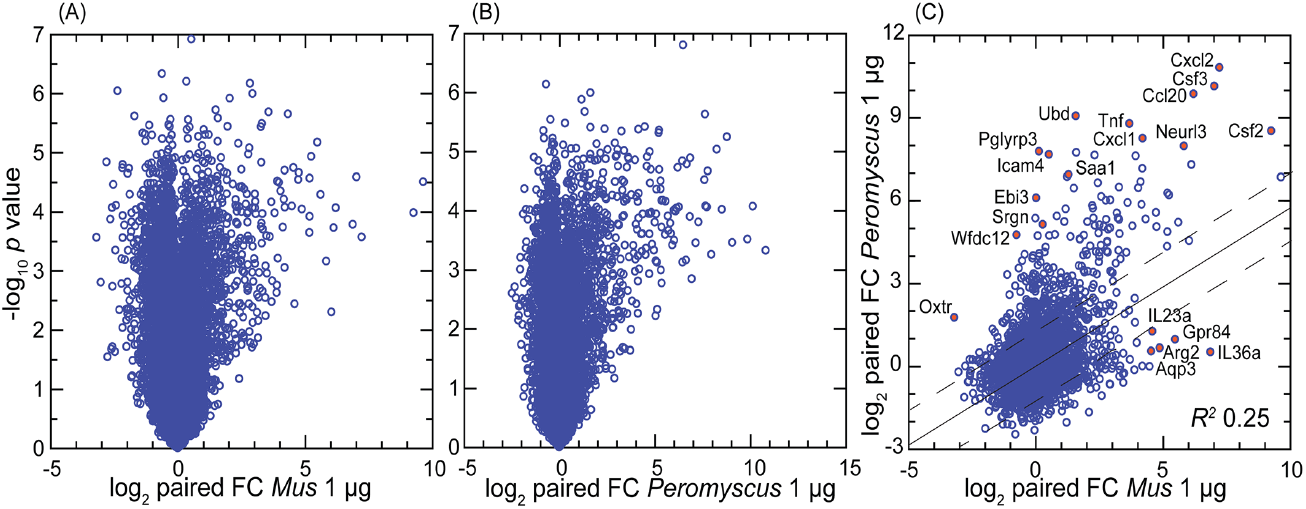
Volcano plot (panels A and B) and scatter plot (panel C) of treatment-to-control fold-change (FC) values from genome-wide bulk RNA-seq of primary dermal fibroblasts of *Mus musculus* (panel A) and *Peromyscus leucopus* (panel B). The treatment was 1 µg/ml of Pam3CSK4 lipopeptide for 4 h. In panels A and B -log_10_-transformed paired *t*-test *p* values are plotted against log_2_-transformed FC for 13,243 *M. musculus* CDS and 12,938 *P. leucopus* CDS with mean TPM ≥10 across all samples. By the criteria of, first, FC ≥2 or ≤0.5 and, second, *p* value <0.001 (FDR <0.0025) there were 149 down-regulated differentially-expressed genes (DEG) and 261 up-regulated DEGs for *M. musculus* (panel A), and 58 down-regulated DEGs and 300 up-regulated DEGs for *P. leucopus* (panel B). In panel C the log_2_-transformed FC values for each species for 14,979 CDS in common are plotted against each other. The linear regression line with 95% confidence interval and coefficient of determination (*R*^*2*^) are shown. Selected CDS that were up-regulated and differentially expressed for each are indicated by name and red fill. Data for analyses are in Dryad Tables D1 (panel B), D2 (panel A), and D3 (panel C).

For the genome-wide RNA-seq for *M. musculus* it was possible that our choice of the top-listed isoform, i.e., instead of all isoforms for the reference set, and which was done for the sake of comparability with the *P. leucopus* reference set, introduced a design bias. To look for evidence of such an effect, we identified 13,786 genes of *M. musculus*, for which there were two isoforms represented in the mouse genome annotation. The sets of corresponding isoforms for same gene were used separately as the references for aligning reads (Dryad Table D7). This analysis identified only a single gene, Gbp6, which encodes the interferon-inducible guanylate binding protein 6, as a DEG with FDR <0.05 that differed between isoform sets. In this case, it was RNA-seq with the second isoform set as the reference that missed this DEG call (Figure S2). From this we concluded that it was not likely that the greater number of isoforms identified for *M. musculus* than for *P. leucopus* substantively biased the analysis one way or the other.

By the gauge of fold-changes of treated over control for paired cultures, the deermouse fibroblasts had a higher number of upregulated DEGs than what was recorded for the mouse cells (Figure 2). Nearly identical results were found at the 10 µg/ml exposure for *M. musculus* (Figure S3), evidence that the lower number of upregulated DEGS in *M. musculus* was not likely attributable to greater resistance of mouse cells to the agonist. By this analysis some genes of relevance to innate immunity that were upregulated in deermice cells but little if at all in mouse cells were Oxtr1 (oxytocin receptor) (32), Ebi3 (Epstein-Barr virus induced gene 3) (33), and the protease inhibitor Wfdc12 (WAP four-disulfide core domain 12), which is homologous to Slpi (34). For *M. musculus* two of the genes specifically upregulated in that species’ cells were Gpr84 (G protein-coupled receptor 84), which is a common loss-of-function allele among inbred strains mouse (35), and the gene for the cytokine interleukin-36A, which is known to be active in skin (36).

By GO term analysis of the DEGs for each species (Figure S4), the responses of the deermouse and mouse cells were largely coherent, with the terms “innate immune response” (GO:0045087), “regulation of inflammatory response” (GO:0050727), and “inflammatory response” (GO:0006954) among the top 5 for each species and with *p* values <10^−10^.

Further differences between the species’ fibroblast cultures were apparent when the DEGs were categorized according to specificity for one species or another, defined here as a ≥10x difference between species in respective fold changes for a given gene (Table 1; Tables S3 and S4). In the most discriminating cases, such as Pla2g2a (phospholipase A2, group IIA) and the PRR Pglyrp3 (peptidoglycan recognition protein 3) (37) of *P. leucopus* or Fth1 (ferritin heavy polypeptide 1) and the siderophore-binding Lcn2 (lipocalin 2) of *M. musculus*, the orthologous gene was not detectably transcribed in the comparison species. For some genes (e.g. Mx2 for *P. leucopus*) the distinctions between species were present under both control and treatment conditions, while for other genes (e.g. Il11 for *M. musculus*) the species difference was only noted for cells exposed to the agonist.

**Table 1.** Differentially-transcribed coding sequences of genes between *Peromyscus leucopus* and *Mus musculus* second passage dermal fibroblasts under conditions of no treatment (control) or treatment with the TLR2 agonist Pam3CSK4 at 1 µg/ml. The left side of table has genes that had higher transcription in *P. leucopus* (P) than *M. musculus* (M), and the table’s right side has genes that had higher transcription in *M. musculus* than *P. leucopus*. The targeted RNA-seq data (Table S2) were normalized by total reads, adjusted for length as reads per kb, and then, for cross-species comparison, the ratio to Gapdh reads for same sample. The 5 different cell cultures for each species (Table S1) were analyzed as pairs within a species for both fold-change (FC) and *t*-test under either control (cont.) conditions or Pam3CSK4 treatment.

| Table 1. |  |  |  |  |  |  |  |  |  |
| --- | --- | --- | --- | --- | --- | --- | --- | --- | --- |
| Higher transcription in *Peromyscus* (P) than *Mus* (M) |  |  |  |  | Higher transcription in *Mus* than *Peromyscus* |  |  |  |  |
| CDS | FC P/M cont. | P/M *t*-test *p* cont. | FC P/M Pam3CSK4 | P/M *t*-test *p* Pam3CSK4 | CDS | FC M/P cont. | M/P *t*-test *p* cont. | FC M/P Pam3CSK4 | M/P *t*-test *p* Pam3CSK4 |
| Arg1 | 14.6 | 6E-04 | 23.2 | 3E-04 | Adamts9 | 2.20 | 2E-01 | 31.3 | 3E-04 |
| Batf2 | 60.2 | 3E-06 | 113 | 4E-06 | Aqp5 | 589 | 8E-09 | 73.9 | 2E-07 |
| Bdkrb1 | 15.5 | 5E-07 | 14.8 | 4E-08 | Arg2 | 5.65 | 2E-03 | 68.7 | 1E-05 |
| Calhm5 | 4.56 | 2E-04 | 13.1 | 6E-07 | Ass1 | 332 | 2E-06 | 1032 | 6E-07 |
| Casp1 | 12.8 | 7E-06 | 26.6 | 1E-06 | Ch25h | 0.66 | 1E-01 | 10.1 | 8E-07 |
| Cd40 | 2.02 | 8E-02 | 13.1 | 7E-06 | Chi3l1 | 80.5 | 2E-06 | 6.24 | 9E-03 |
| Cmpk2 | 8.21 | 1E-03 | 25.8 | 8E-05 | Cxcl11 | 2862 | 2E-08 | 39.9 | 8E-04 |
| Cxcl10 | 5.25 | 9E-04 | 12.6 | 7E-05 | Cxcl5 | 15.7 | 5E-04 | 2.94 | 2E-04 |
| Ddx60 | 13.3 | 5E-04 | 20.5 | 4E-06 | Egln3 | 10.3 | 3E-03 | 11.8 | 4E-03 |
| Dennd3 | 13.7 | 6E-06 | 48.4 | 3E-08 | Ereg | 28.8 | 3E-06 | 21.0 | 8E-05 |
| Deptor | 30.4 | 4E-06 | 27.2 | 6E-06 | Fth1 | 43525 | 1E-09 | 74792 | 4E-10 |
| Dpy19l2 | 14.6 | 4E-07 | 20.0 | 2E-07 | Il11 | 1.92 | 6E-02 | 11.7 | 3E-05 |
| Dtx3l | 5.10 | 2E-06 | 16.2 | 2E-06 | Il1rn | 1530 | 4E-07 | 84.0 | 2E-06 |
| Hpx | 3.35 | 2E-03 | 14.4 | 5E-04 | Il36a | 1.74 | 2E-01 | 142 | 6E-06 |
| Icam4 | 0.96 | 9E-01 | 187 | 3E-10 | Il4r | 12.1 | 2E-06 | 2.70 | 1E-03 |
| Ifi205 | 15.8 | 8E-06 | 41.8 | 1E-06 | Il6 | 17.2 | 1E-05 | 1.75 | 1E-01 |
| Il15 | 2.66 | 2E-03 | 18.6 | 1E-07 | Ism1 | 4.09 | 5E-02 | 12.7 | 6E-04 |
| Irf7 | 4.29 | 4E-03 | 13.2 | 5E-05 | Lcn2 | 1719 | 6E-07 | 12898 | 8E-09 |
| Irgm2 | 5.55 | 5E-05 | 43.7 | 6E-06 | Lrig1 | 4.41 | 2E-04 | 20.8 | 3E-07 |
| Kctd12 | 15.9 | 7E-09 | 17.4 | 2E-08 | Lrrc17 | 90.5 | 3E-06 | 45.6 | 3E-06 |
| Mmp12 | 4372 | 9E-07 | 3114 | 7E-07 | Mmp3 | 11.1 | 4E-03 | 23.4 | 4E-04 |
| Mx2 | 24.3 | 6E-05 | 66.9 | 5E-06 | Mt1 | 18.9 | 1E-06 | 46.4 | 9E-08 |
| Neurl3 | 5.31 | 2E-02 | 21.7 | 2E-04 | Mt2 | 34.6 | 2E-05 | 20.6 | 9E-06 |
| Oas1 | 10.3 | 2E-04 | 24.7 | 8E-06 | Nos2 | 14.9 | 2E-03 | 26.1 | 3E-05 |
| Pglyrp3 | 4.95 | 2E-02 | 1243 | 2E-08 | Rgs16 | 249 | 1E-06 | 640 | 6E-08 |
| Pla2g2a | 882 | 6E-08 | 222063 | 5E-12 | Saa3 | 172 | 2E-04 | 9.81 | 7E-06 |
| Pla2g5 | 58.3 | 9E-07 | 281 | 7E-08 | Sema7a | 5.87 | 5E-03 | 12.0 | 5E-05 |
| Plek | 2.60 | 6E-02 | 18.4 | 8E-06 | Serpina3f | 13.7 | 2E-03 | 3.83 | 1E-02 |
| Rcan1 | 9.90 | 1E-07 | 15.7 | 6E-07 | Serpina3g | 14.1 | 1E-03 | 4.98 | 3E-03 |
| Rgs5 | 375 | 1E-06 | 364 | 2E-06 | Serpina3i | 37.1 | 2E-04 | 38.7 | 2E-04 |
| Rnf122 | 1.68 | 5E-02 | 39.2 | 3E-08 | Slco2a1 | 54.7 | 3E-04 | 123 | 5E-07 |
| Rsad2 | 1.50 | 4E-01 | 23.0 | 7E-04 | Slfn2 | 199 | 7E-11 | 683 | 7E-08 |
| S100a8 | 15.5 | 4E-03 | 29.0 | 4E-05 | Tnfsf11 | 1465 | 3E-06 | 214 | 2E-07 |
| Saa1 | 0.98 | 1E+00 | 57.7 | 7E-07 | Tyk | 7.50 | 5E-04 | 13.6 | 1E-03 |
| Samd11 | 234 | 9E-11 | 97.8 | 3E-08 | Wnt16 | 950 | 1E-07 | 164 | 6E-07 |
| Selp | 1.83 | 1E-01 | 78.3 | 1E-06 |  |  |  |  |  |
| Serpina3n | 128 | 2E-05 | 1742 | 7E-06 |  |  |  |  |  |
| Slc2a6 | 2.99 | 3E-03 | 16.7 | 6E-08 |  |  |  |  |  |
| Slc7a2 | 14.7 | 3E-06 | 3.16 | 5E-03 |  |  |  |  |  |
| Sp110 | 104 | 3E-07 | 59.7 | 4E-08 |  |  |  |  |  |
| Srgn | 0.45 | 4E-01 | 14.4 | 2E-04 |  |  |  |  |  |
| Stat4 | 323 | 2E-06 | 1359 | 5E-08 |  |  |  |  |  |
| Tm4sf19 | 17238 | 9E-12 | 6108 | 7E-09 |  |  |  |  |  |
| Tnf | 0.34 | 5E-03 | 19.3 | 1E-04 |  |  |  |  |  |
| Tnfaip6 | 4.85 | 2E-05 | 20.5 | 6E-08 |  |  |  |  |  |
| Tnfsf18 | 50.3 | 4E-04 | 781 | 2E-06 |  |  |  |  |  |
| Tnfsf4 | 12.3 | 4E-04 | 79.7 | 1E-05 |  |  |  |  |  |
| Traf1 | 2.57 | 3E-02 | 25.4 | 2E-07 |  |  |  |  |  |
| Trim30d | 1649 | 2E-07 | 2293 | 1E-07 |  |  |  |  |  |
| Ubd | 0.87 | 8E-01 | 148 | 1E-05 |  |  |  |  |  |

Many of the genes or pathways that stand out in Figure 2, Table 2, Table S2, or Figure S4 merit further analysis. But to do justice for all is beyond the scope of this report. Accordingly, we limit the focus to the following: genes that prior studies of experimental animals had shown had relevance for the phenomenon of infection tolerance demonstrated by *Peromyscus* species, and DEGs in this study that were unexpected.

**Table 2.**
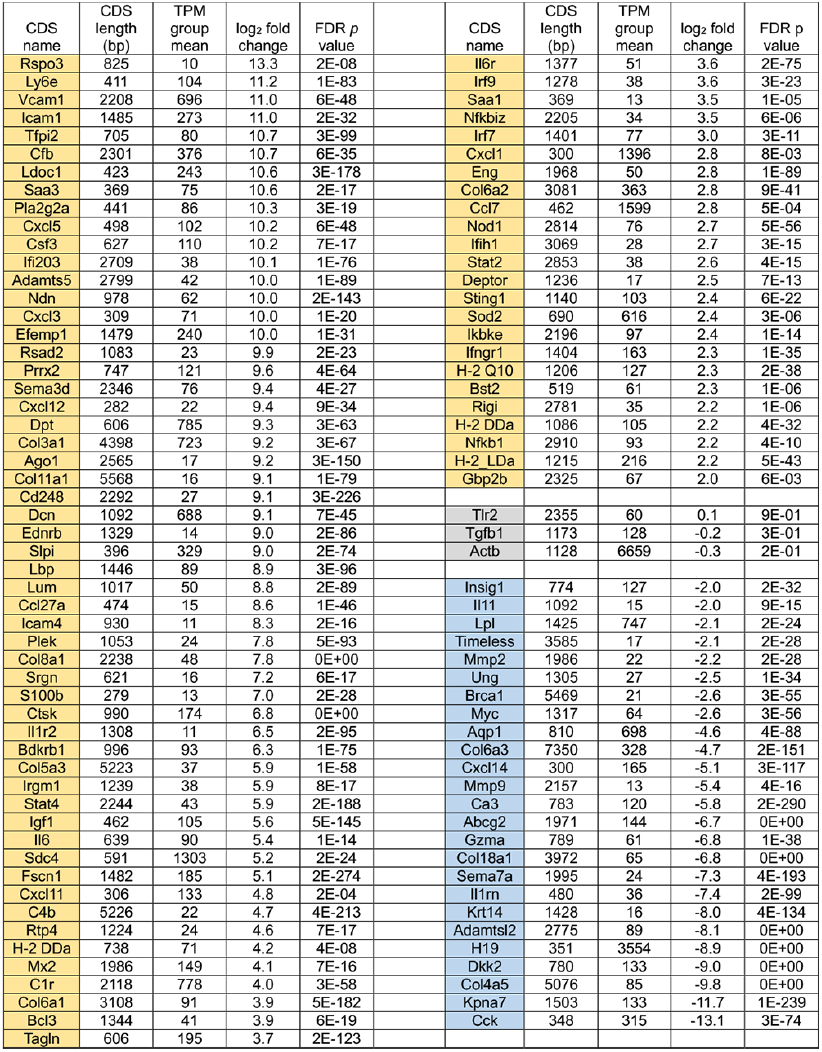
Differentially-transcribed coding sequences (CDS) between low passage (P4) and high passage (P47) dermal fibroblasts of Peromyscus leucopus from bulk RNA-seq analysis of fibroblast cells without or with exposure to Pam3CSK4 or LPS. The comparison was of the samples from 3 conditions for the P4 cells with the samples from the 3 conditions of the P47 cells. The table lists by columns the CDS names, CDS lengths in base pairs (bp), the TPM group mean, the log2 of fold change (FC) between the P4 cells compared to the P47 cells, and the false discovery rate (FDR) p value. The CDS comparatively upregulated in P4 cells are indicated by yellow fill of the cells. The CDS comparatively upregulated in P47 cells are indicated by blue fill. Selected examples of CDS that were not differentially transcribed are indicated by gray fill. The product names of the CDSs are provided in Table S3. The data are drawn from the fuller roster of CDS and results of Table S6.

### Nitric oxide and arginine metabolism pathways

A notable finding in prior studies of *P. leucopus* and *M. musculus* animals in response to LPS was a dichotomy between species in the blood and spleen in relationship between expression of nitric oxide synthase 2 (Nos2) and arginase 1 (Arg1) (19, 20). Deermice under treatment displayed a high Arg1-to-Nos2 ratio, which was consistent with the profile for alternatively activated (or M2) macrophages. The mice instead had an inverted ratio of Arg1-to-Nos2, which is more typical for classically activated macrophage (or M1) polarization (38, 39). The demonstration that LPS-treated, low passage fibroblasts of P. leucopus did manifest transcription of Nos2 above baseline values indicated that this species had the capacity for expressing Nos2.

In the present study we confirmed the transcription in *P. leucopus* cells of Nos2 by RNA-seq with different set of dermal fibroblast cultures and with a different TLR agonist (Figure 3 and Table S3). The fold-change increase in Gapdh-normalized transcription of Nos2 between control and agonist-treated cells was similar in both species, but the expression of Nos2 was several fold lower at baseline in the deermouse cells. This was accompanied by a high baseline transcription of Arg1 with modest further elevation with agonist exposure for the *P. leucopus* cells. While the *M. musculus* fibroblasts, like its cells in the blood, had low transcription of Arg1 under control and treatment conditions, transcription of Arg2, the gene for the mitochondrion-based type II arginase (40), was many fold higher than observed for deermouse cells for both conditions (Figures 2 and 3; Table 1).

**Figure 3.**
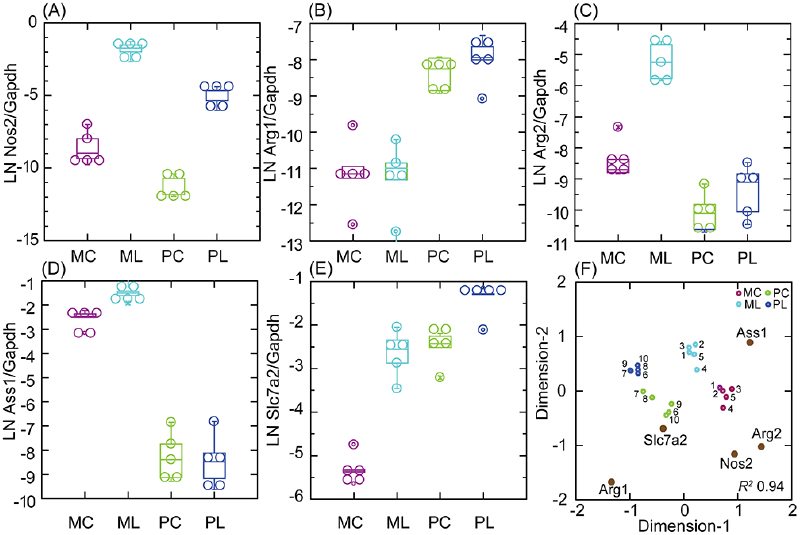
Transcription of nitric oxide and arginine metabolism CDS Nos2, Arg1, Arg2, Ass1, and Slc7a2 in targeted RNA-seq of primary dermal fibroblasts of *M. musculus* (M) or *P. leucopus* (P) without (C) or with (L) treatment with the lipopeptide Pam3CSK4 at 1 µg/ml. Panels (A)-(E) are box-whisker plots of individual CDS. Panel (F) is a Multi-Dimensional Scaling configuration plot of a analysis reduced to 2 dimensions (*x*- and *y*-axes) from the data for the box plots and with coordinates of CDS variables for the 10 samples indicated by gene names and symbols with dark brown fill. Pairs of individual lineages (1-5 for P and 6-10 for M; Table S1) are designated by numbers. The data for analyses are in Table S2.

Of note were other arginine metabolism genes that in their profiles of transcription were distinguishing between the species (Figure 3 and Table 1). One of these was Ass1, the gene for arginosuccinate synthetase 1, which also is based in mitochondria (41). Ass1 was little transcribed by *P. leucopus* cells but highly at baseline and further more with treatment in *M. musculus* cells. A second was Slc7a2, which encodes a cationic amino acid transporter for L-arginine (42). Like Arg2 in the mouse cells, Slc7a2 transcription increased to high level in agonist-treated cells. Unlike Arg2, which was little transcribed in *P. leucopus*, Slc7a2, was at baseline at the same normalized level as for the treated *M. musculus*. Taken together, Nos2, Arg1, Arg2, Ass1, and Slc7a2, could in their multi-gene profiles differentiate not only between species but also between conditions in each species (Figure 3 panel F). The findings also point to differences between in the metabolism of arginine and the extent to which this takes place in mitochondria of cultured fibroblasts.

### Secretory leukocyte peptidase inhibitor (Slpi)

Another feature distinguishing *P. leucopus* from *M. musculus* after exposure to LPS was a more than thousand-fold increase over baseline of transcription of Slpi in blood, as documented by both RNA-seq and RT-qPCR. At the same time there was only marginal elevation in the treated mouse samples over the low levels observed for controls (19). A heightened expression of Slpi over baseline was also seen in *P. leucopus* infected with the relapsing fever agent *Borrelia hermsii*, as well as in a second experiment with LPS and with outbred mice (20). For LPS-treated cultures of ear skin fibroblasts, comparatively high transcription of Slpi was noted not just in the treated cells but in untreated cells as well (19).

In the present study with an independent set of cultures from *P. leucopus* and which were processed in parallel with cultures from outbred *M. musculus*, we confirmed high transcription Slpi in untreated cells and, as before, with a modest increase after TLR2 agonist exposure (Table S3). For this study we also documented the expression in the *P. leucopus* fibroblasts of Slpi protein by MALDI-TOF. The following three peptides, which cover 46% of the 106 amino acid processed protein (accession XP_028724460.1) were identified: KDSIKIGACPSISPAK, CTVPLPISRPVRRK, and KSGKCPTFQGRCMMLNPPNK.

The various pathways leading to Slpi expression are not well defined. To provide insight for this we identified among 365 genes, which were upregulated DEGs for either or both *P. leucopus* and *M. musculus* (Table S2), those that were highly correlated with Slpi transcription across species and conditions. The first transcription factor on the list of descending *R*^*2*^ values was Nfe2l2, more commonly known as Nrf2, which encodes the protein nuclear factor, erythroid derived 2, like 2. Nfe2l2 is a transcriptional regulator of a number of genes involved in the adaptive response to oxidative and other cytotoxic stress (43).

Nfe2l2 transcription in the deermouse and mouse fibroblasts was compared with that of Nfkb1, the gene for a more broadly active transcription factor (Figure 4). For the control cells, when Nfkb1 transcription was about the same in both species, Nfe2l2 expression was nearly as high for *P. leucopus* as it would be after exposure to the lipopeptide. While Nfe2l2 transcription in the treated *M. musculus* cells never reached the levels observed in *P. leucopus*, there was a still greater fold increase over baseline than was observed for deermouse cells. In contrast, the relationship between Nfe2l2 and Slpi was more direct and consistent across species and conditions, suggesting a regulatory role of Nfe2l2 in Slpi expression in *P. leucopus*.

**Figure 4.**
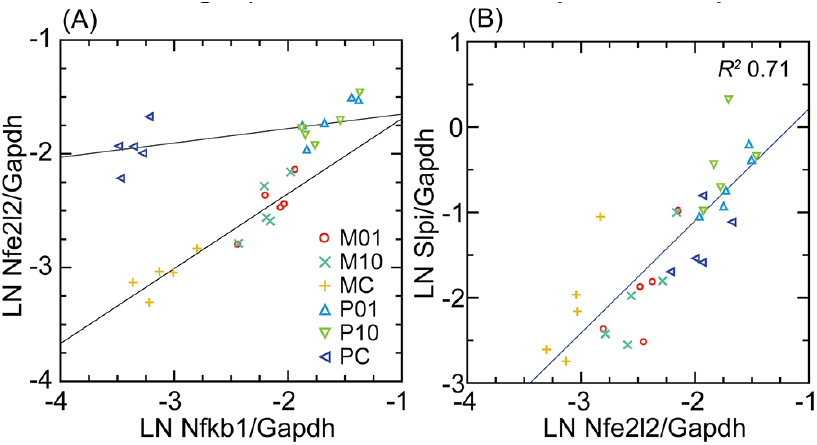
Scatter plots and linear regressions of normalized transcription of either Nfe2l2 (Nrf2) on Nfkb1 (panel A) or Slpi on Nfe2l2 (panel B) for *M. musculus* (M) and *P. leucopus* (P) dermal fibroblasts that either untreated (C) or treated with Pam3CSK4 lipopeptide at 1 µg/ml (01) or 10 µg/ml (10). For analysis of panel (A) the regression are separate for P and M. In panel (B) the regression was for both species and with the coefficient of determination (R2) for both. Data for analyses are in Table S2.

### PRRs and interferon-stimulated genes (ISG)

The results with the fibroblast cultures replicated some of the previous findings with experimental animals. One observation for those PRRs and ISGs that were DEGs for either or both mouse and deermouse was that illustrated by the genes for the PRR RIG-I and Isg15 in panel (A) and panel of Figure 5, namely transcription that was lower in *P. leucopus* than *M. musculus* at baseline but then exceeded that of the mouse with agonist exposure. Another observed pattern is shown in panel (C). As for *P. leucopus* animals, the antiviral ISG Mx2 in the fibroblast cells was at an elevated level relative to mouse at baseline and displayed an even higher level with treatment. Other genes demonstrating the second pattern in the fibroblasts, replicating the findings for the experimental animals, were the PRR Ifih1 (MDA5), the regulatory factor Irf7, and the antiviral ISG Oas1 (Figure S5).

**Figure 5.**
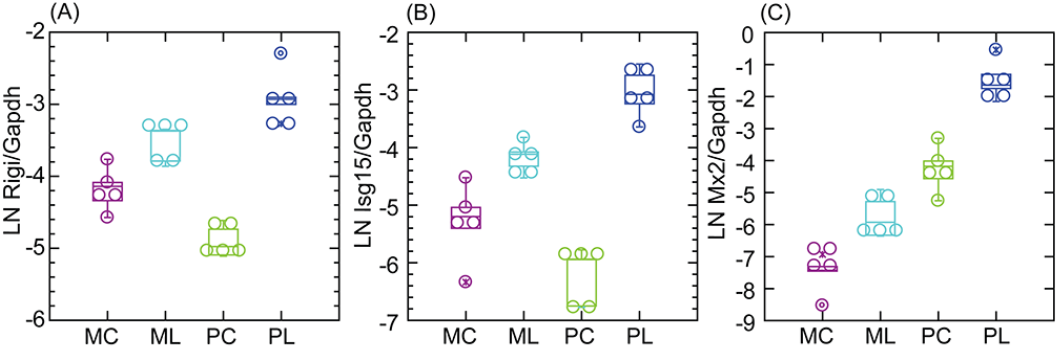
Box-whisker plots of log-transformed Gapdh-normalized transcription of the Pattern Recognition Receptor (PRR) CDS Rigi (panel A) or the Interferon Stimulated Genes (ISG) Isg15 (panel B) or Mx2 (panel C) in *M. musculus* (M) or *P. leucopus* (P) dermal fibroblasts without (C) or with treatment (L) with lipopeptide Pam3CSK4 at 1 µg/ml. Data for analyses are in Table S2.

### Endogenous retrovirus and transposable elements

In a previous analysis of blood of deermice and mice, we observed that the two species differed in their transcription of sequences for a retroviral envelope (Env) protein and Gag-pol polyprotein in response to the TLR4 agonist LPS (20). For the present study we used much expanded reference sets of ∼1 million annotated ERV and derived transposable elements (TE) sequences for each species (Dryad Tables D4 and D5). The distribution of lengths of the ERVs in the complete sets was remarkably similar between species, with approximately equal proportions of sequences in different size classes, including those more the 5 kb in length (Figure S6 panels A and B). After exclusion of ∼90% of the sequences with lengths under 500 bp, there remained 104,932 sequences for the *M. musculus* reference set and 103,397 for *P. leucopus* (Dryad Table D6). For these sets the lengths ranged from 500 to 9329 bp for *M. musculus* and from 500 to 9543 bp for *P. leucopus*. The distributions in sequence lengths were skewed toward lower values and with a long tail to the right; the skewness statistic was 4.3 for mouse and 3.5 for deermouse. The median length (inter-quartile range) was 711 (572-950) bp for M. musculus and 742 (567-1164) bp for *P. leucopus* (Dryad Tables D8 and D9). For the control samples, mean values for the TPM-based transcription measure were 9.6 for *M. musculus* and 9.1 for *P. leucopus* sequences.

A closer look at the ERV loci with higher transcription levels in cells across all conditions revealed differences between species. If the sets of sequences were limited to the top 1000 in descending order of mean TPM (i.e. the top ∼1% for each species), the species were comparable in terms of the distributions of the length-adjusted TPM values (Figure S6 panel C). What distinguished the mouse sequences were their overall longer absolute lengths (Figure S6 panel D). The mean lengths were 2622 (2525-2719) bp for mouse and 985 (942-1029) bp for deermouse (*t*-test *p* <10^−10^). The lengths difference was as marked for the top 100 (i.e. ∼0.1%) of transcribed sequences by TPM values; mean lengths for the top 0.1% were 3019 (2742-3186) bp for mouse and 1146 (972-1319) for deermouse cultures.

We compared the 5 fibroblast lines each for *P. leucopus* and *M. musculus* for the fold-change differences in transcription between low-passage cells treated with 1 µg/ml Pam3CSK4 and untreated cells. As we observed for the genome-wide CDSs (Figure 2), the numbers of DEGs with higher transcription after agonist exposure were greater than the number of DEGs with reduced transcription post-exposure for both species (Figure S6 panels E and F). There were sub-sets of 87 ERV/TEs each for *P. leucopus* and *M. musculus* that increased ≥2-fold in transcription in the presence of 1 µg/ml lipopeptide and for which the paired *t*-test *p* value was <0.001 (FDR <0.05) (Table S4). For cross-species comparisons we used *Z* scores for lengths, which were based on the distributions of lengths for the complete reference sets of ∼100,000 sequences for each species, as well as Z scores for mean TPM across all conditions, based on distributions 5296 mouse and 7432 deermouse ERV/TEs with mean TPM ≥10 (Figure 6). Between these sub-sets the mouse ERV/TEs tended to have higher mean transcription, but there was considerable overlap between species in the ranges for this measure (Figure 6 panel A).

**Figure 6.**
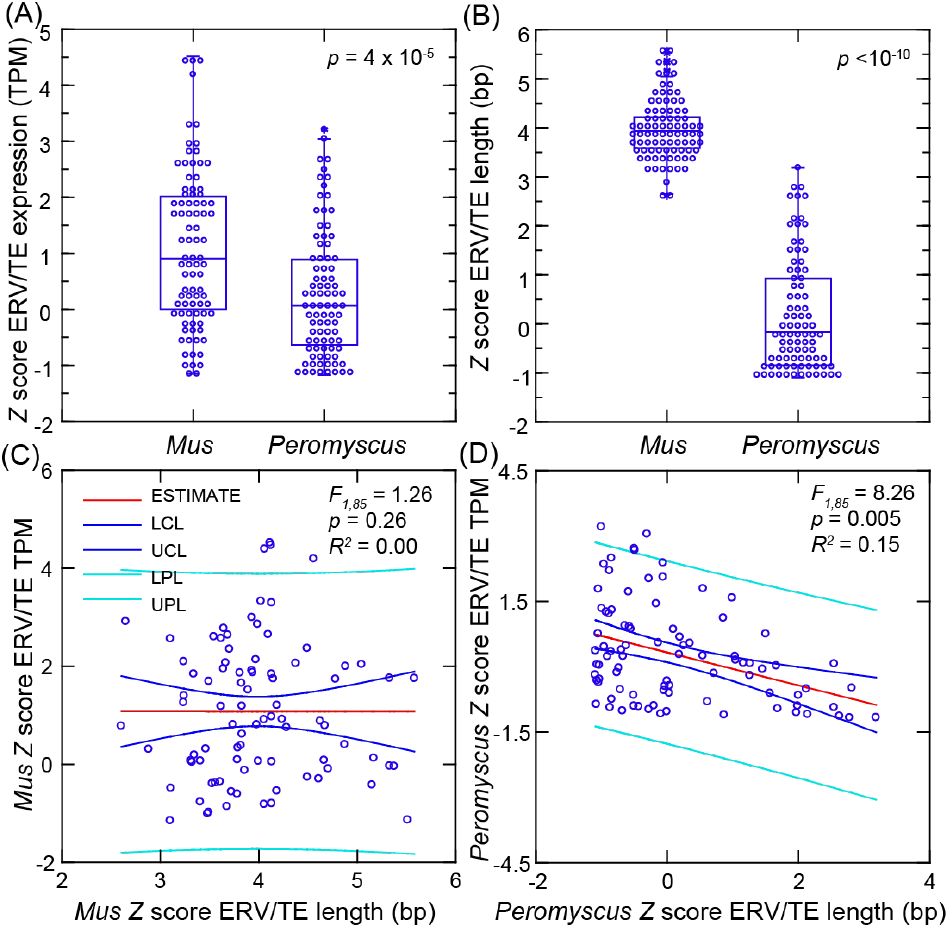
Comparison of the differentially-expressed genes (DEG) for ERV/TEs in five low-passage dermal fibroblast cultures each for *Mus musculus* and *Peromyscus leucopus* in the presence or absence the TLR2 agonist Pam3CSK4. Panel A is a box-whisker plot of *Z* scores of the mean TPM across the samples for each species for the 87 DEGs of *M. musculus* and 87 DEGs of *P. leucopus*. Panel B is a box-whisker plot of the *Z* scores for lengths of the corresponding DEGs. The non-parametric test is Kruskal-Wallis for the *p* values of panels A and B. Panels C and D are General Linear Model regressions with estimates (mean), lower control limits (LCL), upper control limits (UCL), lower prediction limits (LPL), and upper prediction limits (UPL) for ERV/TE mean TPM on length for the DEGs of *M. musculus* (panel C) and DEGs of *P. leucopus* (panel D). Data for analyses are from Table S4.

The characteristic more definitively distinguishing the species for these DEG sub-sets was length (Figure 6 panel B). These *Z* scores corresponded with ranges of lengths of 2479-8726 bp for *M. musculus* and 500-4162 bp for *P. leucopus* (Table S4). Not only were *P. leucopus* DEGs shorter in general than the mouse DEGs, but there was a trend of lower transcription with increasing length of the deermouse sequences that was not observed in mouse sequences (Figure 6 panels C and D).

While there were no *P. leucopus* sequences in this sub-set longer than 4162 bp, let alone 5000 bp, 21 (24%) of 87 *M. musculus* up-regulated ERV/TEs were ≥5000 bp (*p* < 10^−10^). We identified in these 21 mouse sequences ORFs having an ATG as start codon (where A is position +1), were at least 30 codons, and had both a purine at position -3 and a G at position +4. These are features of a Kozak sequence for a protein translation initiation site in vertebrates (44). They identify ORFs that more plausibly are translated than ORFs lacking these features. Of these 21 DEG ERV/TEs, 19 (90%) collectively had 115 ORFs meeting this criterion on either plus or minus strand and ranging in length from 30 to 1091 amino acids and a mean of 92 (67-116) (Table S5). Of these, 46 (39%) would encode all or part of an ERV Gag-pol polyprotein (n=45) or Env protein (n=1), in total constituting a mean proportion of 0.22 (range 0.06-0.44) of the lengths for 15 qualifying ERV/TEs. In contrast, for the 6 longest *P. leucopus* DEG ERV/TEs, which ranged from 3034 to 4162 bp (Table S5), there were only 12 short ORFs (range of 93-258 bp) that met the Kozak sequence criteria, and only 1 of the predicted peptides of 31-86 amino acids had discernible similarities to either Gag-pol polyproteins or Env proteins of ERVs.

In sum, the mouse ERV/TEs that were up-regulated in transcription in the dermal fibroblasts exposed to the TLR2 agonist were not only substantially longer than the DEG sequences for the deermouse fibroblasts, but there was also evident greater potential for translation of whole or parts of ERV/TE proteins than was noted for the deermouse DEGs among the ERV/TEs.

### Interleukin-11

Unlike preceding examples, which were of correspondences between the findings of in vivo and in vitro systems, some distinguishing genes for the fibroblasts would not have been predicted by studies of blood, spleen, or liver. One we highlight here is interleukin-11 (IL-11), the gene for which was an upregulated DEG in mouse fibroblasts, but lowly transcribed in deermouse fibroblasts, regardless of condition (Table 1). IL-11 is a member of IL-6-type cytokine family; the specific receptor, IL-11Ra, is expressed by Il11ra in fibroblasts but not immune cells (45). An upstream transcription factor for IL-11 is NF-kB (Nfkb1). IL-11 is considered a key factor in the inflammation of aging (“inflammaging”) process, in part by promoting fibrosis (46, 47).

A targeted RNA-seq analysis of transcription of Nfkb1, Il6, Il11, and Il11ra, with Gapdh as the housekeeping gene for normalization across species, is shown in Figure 7. There was higher baseline levels of Il6 in mouse cells than deermouse cells, a greater magnitude of Il6 elevation with agonist exposure in deermouse cells, and similar Nfkb1 levels between species. The finding of the genome-wide RNA-seq analysis for Il11 was confirmed. While Il11 transcription levels increased in mouse cells with both concentrations of Pam3CYSK4, in deermouse there was, if anything, a further decline from low levels at baseline. The scarce to absent Il11 transcripts in these low-passage *P. leucopus* cells was not accompanied by reduced expression of its receptor Il11ra, which was at comparatively high levels in deermouse control cells and, unlike mouse cells, did not decline with agonist exposure. These were the findings for the early passage cells of *P. leucopus*. As described below for high passage cells, the capacity of *P. leucopus* for transcription of Il11 had not been lost.

**Figure 7.**
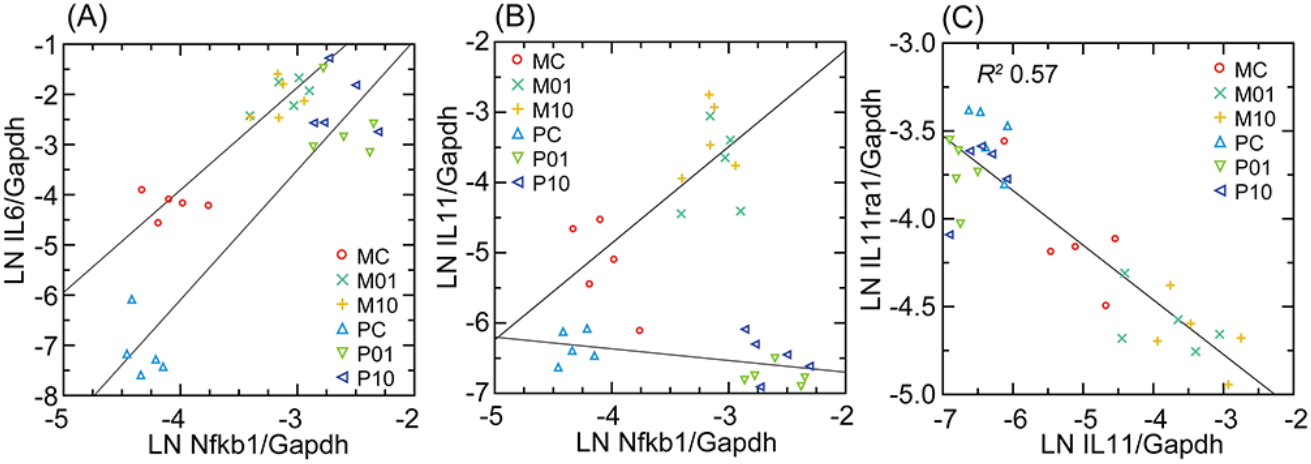
Interleukin-6 (Il6; panel A), interleukin-11 (Il11; panel B), and interleukin-11 receptor (Il11ra; panel C) Gapdh-normalized transcription compared in scatter plots and with linear regressions for *M. musculus* (M) and *P. leucopus* (P) dermal fibroblasts that were either untreated or treated with Pam3CSK4 lipopeptide at 1 µg/ml (01) or 10 µg/ml (10). For analyses of panels (A) and (B) linear regressions are separate by species. For panel C the linear regression with coefficient of determination (*R*^*2*^) was for both. Data for analyses are in Table S2.

### Single-cell RNA-seq of low passage populations

At their outsets, the cultures of the full-thickness ear tissue included cartilage, capillaries and other small vessels, epidermis, hair, and hair follicles as well as the dermis that would be the presumptive source for the fibroblasts (48). What emerged from this initial mixture of cell types were adherent cells for both *P. leucopus* and *M. musculus*. These could be released from the polystyrene dish bottoms with trypsin, and when these were used to populate a fresh culture, the cells soon became adherent again. They had the morphology of fibroblasts (Figure 1), and we had the bulk RNA-seq results, which for both species, confirmed the expression of fibroblast markers, such as the collagen gene Col1a1, fibroblast-specific protein 1 (S100a4), and vimentin (Vim). But heterogeneity was undefined.

As a first step to better characterize these primary cultures we carried out single cell RNA-seq of two sets of pooled cells, one for each species. Each pool contained equal parts of cells that were untreated, cells exposed to Pam3CSK4 for 4 h, and cells exposed to LPS for 4 h. The reasoning was to represent in a single population cells after exposure to two different TLR agonists as well as cells at baseline and then processed together. Each species’ sample yielded ∼4 × 10^4^ cells and, after downstream processing, indexing, library preparation, and sequencing, ∼10^9^ PE150 reads.

We approached the analysis without set ideas about markers to prioritize. Gene expression matrixes were normalized and dimensionally reduced by principal component analysis. UMAP projections and cell clustering were generated by the top principal components for each cell. There were 10 major clusters for each species (Figure 8 panel A), numbered 1-10 for *P. leucopus* and 11-20 for *M. musculus*. For both species’ cells these organized in two super clusters, shown on the left and right in each graphic. The individual clusters for each species were in roughly the same spatial arrangement, with the exceptions of cluster 10 of *P. leucopus*, and cluster 20 of *M. musculus*, which did not have clear counterparts in the comparison species. By analysis of single genes we found that transcription of two collagen genes, Col15a1 and Col8a1, specifically distinguished left and right super clusters, for both species, thus providing evidence of comparability of deermouse and mouse cultures in terms of cell type representation at this higher order.

**Figure 8.**
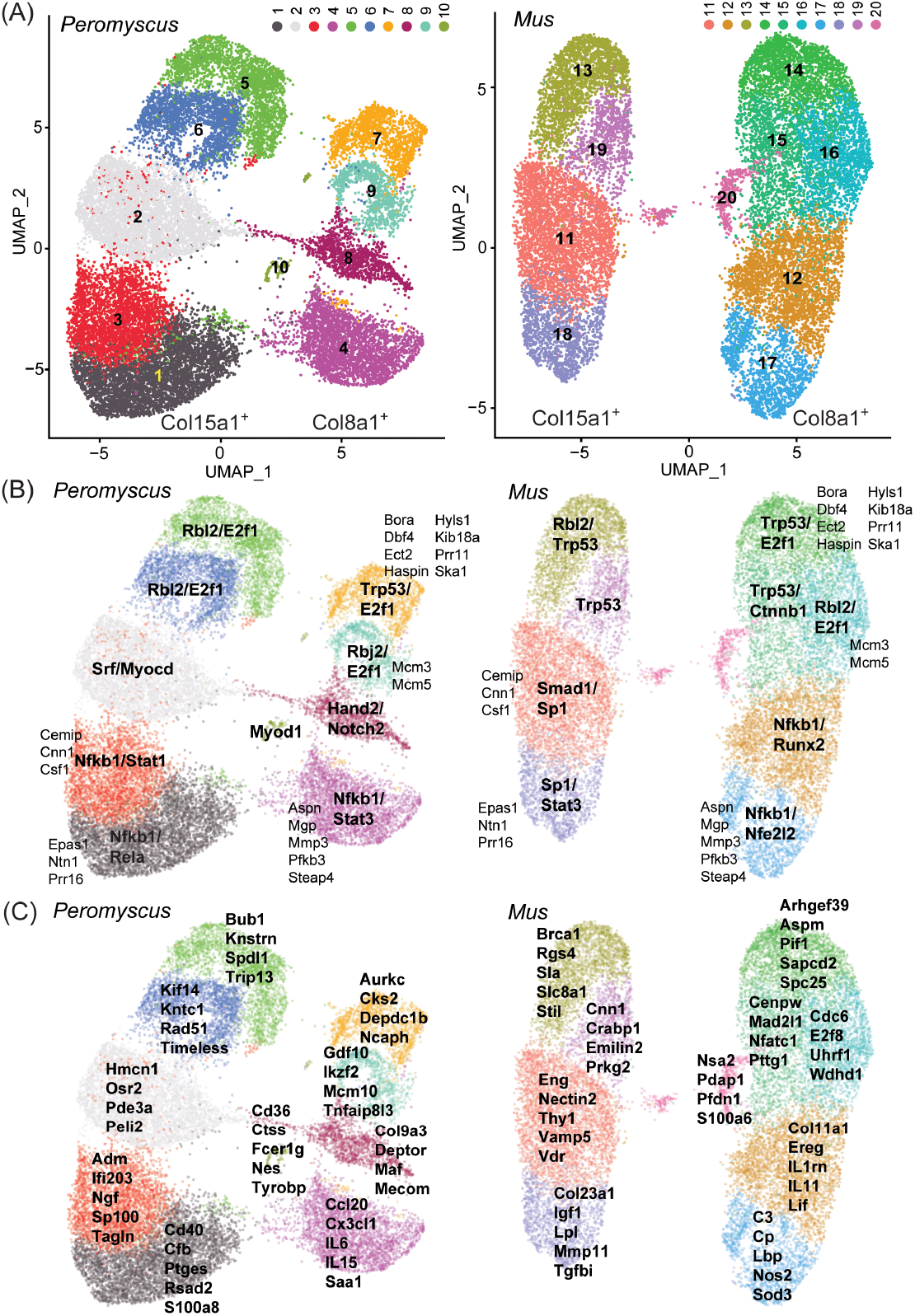
Single cell RNA-seq of low passage cultivated dermal fibroblasts of Peromyscus leucopus and Mus musculus with 2-dimensional UMAP projections of clusters of types of cells in pooled populations of cells that were untreated or exposed to either Pam3CSK4 or LPS. In panel (A) the clusters of each species are identified by number and color: 1-10 for P. leucopus and 11-20 for M. musculus. Two CDS that served as markers for the left and right super clusters were Col15a1 (left) and Col8a1 (right). In panel (B) transcription factors that were predicted by GO term analysis to be associated with signifying CDS for each cluster (Table S5) are indicated in bold font. CDS that defined the same clusters in both species are in smaller and regular font. In panel (C) CDS that were enriched in representation in the clusters and that discriminated to that degree between species are listed for each cluster.

Provisional cross-species analogies among clusters were supported by predictions of what transcription factors would be predominant from GO term analysis of the genes contributing to cluster discrimination (Figure 8 panel B and Table S6). Transcription factors associated more generally with normal cell growth, such E2f1 and Rbl2, appeared to define the upper clusters (i.e. 5, 6, 7, and 9 of *P. leucopus* and 13-16 and 19 of *M. musculus*) of each of the left and right super clusters of deermouse and mouse. On the other hand, the lower poles of both super clusters, were associated with transcription factors, such as Nfkb1, Nfe2l2 (Nrf2), Stat1, and Stat3, more typically identified with host responses to a stressor, suggesting to us that these lower pole clusters were the cells exposed to one or the other TLR agonist. For half the clusters there were also between 2 and 8 genes that contributed to defining a cluster in the same relative location in both deermouse and mouse. These were clusters 1, 3, 4, 7, and 9 for *P. leucopus* and 11, 14, and 16-18 for *M. musculus*. These included the Epas1 (endothelial PAS domain protein 1) for clusters 3 and 11, and Csf1 (colony stimulating factor 1, macrophage) for clusters 1 and 11.

If only the 4-5 top-ranked genes by degree of enrichment and that uniquely contributed to cluster identity were considered (Figure 7 panel C), we observed again in the lower halves of the two super clusters a predominance of genes that are associated with innate immune and other host responses. Of note with regard to other aspects of this study, are the anti-viral effector Rsad2 and calgranulin component S100a8 in cluster 1, interleukin-6 and the chemokine Cxcl20 in cluster 4, interleukin-11 and the interleukin 1 receptor antagonist Il1rn in cluster 11, and Nos2 and complement component C3 in cluster 17. The small cluster 10 located between the two super clusters of *P. leucopus* stood out from the other 9 clusters of the deermouse fibroblasts and all 10 mouse clusters in the specifying contributions of dendritic cell-associated genes Cd36, Fcer1g (CD23), and Tyrobp (DAP12). Cluster 20 of *M. musculus* was defined in part by the genes listed in the panel C of figure but also by enrichment of lower magnitude for several ribosomal proteins (Table S6), an indication that this was a more rapidly dividing cell population, without a counterpart in the *P. leucopus* fibroblast sample.

### Spontaneously transformed *P*. *leucopus* fibroblasts

Bulk RNA-seq was carried out on cultures of *P. leucopus* at passage 4 (P4) and passage 47 (P47) and which were controls or had been treated with Pam3CSK4 or LPS for 4 h. The number of PE150 reads ranged between 2.13-2.93 × 10^8^ with a mean of 2.45 × 10^8^. The primary objective was to identify genes that were transcribed in P4 cells but to little or no degree in P47 cells or vice versa. For this purpose we compared the 3 samples of P4 against the 3 samples of P47 cells across conditions. By the criterion of a fold-change of ≥4, mean TPM ≥10 across all samples for a given gene, and an FDR *p* value <0.01, 80 genes were differentially upregulated in P4 cells and 25 were upregulated (denoted by negative fold-change values) in P47 cells (Table 2 and Table S7). Genes that were little transcribed in P47 compared to P4 cells were Col8a1, Deptor, Eng, Il6, and Lbp, each of which were more closely associated with the right-side super clusters in the single cell experiment (Figure 8). On the other hand, upregulated genes in the transformed cells included Il1rn, Il11, and Timeless, which were associated with the left super cluster. This indicated that the transformed cells more likely trace their lineage to the left super cluster fibroblasts than the fibroblasts on the right.

Besides the near-complete or complete loss of expression of several of the innate immunity-associated transcription factors, cytokines, chemokines, and ISGs that featured in the comparison with low-passage mouse fibroblasts, there was also diminished transcription of MHC class I protein genes in the P47 cells. In addition to the upregulated Il11 and Il1rn, which notably were lowly transcribed in early passage deermouse cells in comparison to mouse (Table 1), two cancer-associated genes, Brca1 and Myc, were more highly transcribed in P47 than in P4 cells.

A secondary objective was to identify in the transformed cell line those genes that not only were still transcribed but also increased in expression after exposure to either or both of the TLR agonists (Table S8). If this was the case, it would indicate retention of some signaling pathways of cell autonomous immunity. One of these was Slpi, which was a thousand-fold lower in transcription in P47 cells than the P4 cells across all conditions (Table 2), but still was ten-fold higher in transcription in the agonist-treated P47 cells than in the controls. Another was Cxcl1, which was a hundred-fold higher in transcription in Pam3CSK4-treated P47 cells than in the controls. The chemokine Cxcl10 was not one of those diminished in transcription in P47 versus P4 cells, but, like Cxcl1, it displayed greater than ten-fold up-regulation in transcription with the TLR2 agonist but not the TLR4 agonist.

The reads were also aligned with *P. leucopus* reference set of 103,397 ERVs of ≥500 bp. Across the 6 samples, 10,843 (10.5%) ERV/TEs had a mean TPM ≥ 5. Of these, 832 (7.7%) of ERV/TE sequences were more highly transcribed in low-passage than in high-passage cells across all 3 conditions by the criterion of FDR *p* <0.01 and absolute fold-change ≥2 for 3 samples for each passage history (Table S9). By the same criteria, 1136 (10.5%) were more highly transcribed in high-passage cells (denoted in the table as negative fold-change values).

The distinguishing DEGs of P47 cells were of interest because of their association with an ostensibly immortalized state. The high passage cells’ upregulated DEGs were in general longer (mean of 1189 vs. 1082 bp; *p* = 0.01) and more highly transcribed (mean TPM of 328 vs. 81; *p* = 0.003) than low passage cell DEGs. Of the 14 high passage cell DEGs of ≥5 kb size, 3 were >9 kb, with lengths (and chromosome locations) of 9543 (NC_051064.1:6812577-6822120), 9432 (NC_051076.1:33708849-33718281), and 9208 (NC_051063.1:54636550-54645758) bp. Not only do the three have ORFs for components of Gag-pol polyproteins, including reverse transcriptases, but they also would encode retroviral envelope (Env) proteins. For the ERVs of 9543 and 9432 bp, the CDS for Env sequences aligned with feline leukemia virus (FLV) proteins along their full-lengths and without frame shifts or in-frame stop codons. The Env CDSs of the 9543 bp and 9432 bp ERV sequences have a purine at -3 position and ATGG for positions +1-4, thus plausibly translatable. The 1983 nt sequence of the predominant FLV-related Env polyprotein mRNA (GenBank accession PV892740) transcribed by the transformed *P. leucopus* fibroblasts would encode a polypeptide of 661 aa with a gp70 portion from positions 32 to 463 and the more conserved, transmembrane p15E portion from positions 464 to 638.

In contrast, the 6 DEGs of ≥5 kb, fold-change ≥2, and FDR *p* <0.01 of low passage cells were of a maximum length of 7503 bp (Table S9). The 6 ERV sequences each comprised ORFs for parts of Gag-pol polyproteins, but none would discernibly encode a retroviral envelope protein, in part or whole. This and other findings about the immortalized fibroblasts showed that amidst the silence of many genes of inflammation consequence, such as Il6, other genes, such as some ERV sequences and Il11, were effectively released from repression.

## Discussion

Previous comparisons of *P. leucopus* with *M. musculus* used laboratory-bred animals that were euthanized at experiment’s end (19, 20, 49, 50). Acknowledging that in vivo studies deserve their pre-eminence in research, we asked whether pre-mortem primary cultures of a tissue from *P. leucopus* could stand-in for the animals themselves for some research questions. For field-based studies this would allow for the release of captured animals after measurements and specimens have been taken. This was one rationale for the application of wing tissue biopsies for studies of bats in nature (51, 52). For laboratory-based investigations of stock colonies longitudinal designs with non-lethal sampling of tissue at different times become feasible.

An indisputable advantage of tissue culture is the capability to freeze the cells indefinitely, permitting replication of an experiment or the application of procedures not available at the time of collections. With their tissues propagated and cultures split, animals can serve as their own controls in experiments. There can be interventions that could not be feasibly tested with live animals, because of toxicity at the organismal level or impeded delivery to the tissue or organ of interest. Experiments on cultured cells do not suffer from these restrictions. These manipulations include RNAi and other non-transgenic gene silencing methods or CRISPR gene editing applications before this technology is achieved for the whole animal.

These are justifications for use of cultured cells. But their validity as approximations of live animals rests on two assumptions. The first is that cells cultured from a non-model organism, e.g. *P. leucopus*, sufficiently resemble those of a more established model organism, i.e. the house mouse, that one can exploit extant databases and literature on the model organisms. If there are many incongruities or misalignments, the findings for the deermouse might be limited in impact to that one genus. The second assumption is that in vitro experiments’ results generally line up with those in vivo experiments for the same organism. In other words, if a gene is upregulated in the animal under a certain condition, is it also upregulated in the tissue culture cells?

In considering the first assumption, we noted the report that established cultures of *P. leucopus* fibroblasts had similar doubling times to those of *M. musculus* (53). Unlike cultured fibroblasts of humans and some other large mammals, deermouse and mouse cells had telomerase activity and did not undergo replicative senescence in the Seluanov et al. study (53). The observed decline in the present study of one *M. musculus* culture within a few passages is provisionally attributed to greater oxygen sensitivity of the mouse cells (29) and not replicative senescence. Without discounting a possible distinction between species in their responses to oxidative stress, the early passage populations of *P. leucopus* and *M. musculus* dermal fibroblasts had more in common than differences by genome-wide RNA-seq (Figure S1; Figures 2 and 3).

With acknowledgement that analogies between scRNA-seq UMAP clusters for different species should be made with caution, we propose that assignment of correspondences between species was justified not only on the basis of their locations in the maps but also signifying genes in common (Figure 7; Table S6). Col15a1 and Col8a1 empirically distinguished the left and right super clusters of both deermouse and mouse in this experiment (Figure 7), but these genes are not generally recognized as markers for the predominant types of fibroblasts in mouse skin: papillary fibroblasts in the upper dermis and reticular fibroblasts in the lower dermis (54). Some markers known for their specificities for fibroblast types in intact skin would be expected to decline in expression in culture (55), thus reducing the numbers of informative characteristics available for cell origin typing. Nevertheless, there were some enrichments of expression that allow for tentative assignment of origins for some clusters. Examples were Crabp1 (cellular retinoic acid binding protein 1) and Col23a1 for left sided clusters and Aspn (asporin) and Col11a1 for right-sided clusters (Table S6). These suggest to us that the left and right super clusters had origins in papillary fibroblasts and reticular fibroblasts of the dermis, respectively, of mouse and deermouse.

Justifying the second assumption is a challenge, what with differences in complexity and time scale between the in vivo and in vitro. But, as noted by Tyshkovskiy et al. (56), who cited examples from research on aging, a rationale for this assumption exists. One of those examples was the report of Ma et al. (57). These investigators cultivated fibroblasts from the skin of captured *P. leucopus* and *P. maniculatus*, along with several other small and medium sized mammals, and found that the RNA-seq and biochemical assay results with in vitro-cultivated cells generally conformed with known longevity traits of the animals themselves. We similarly noted correspondences between prior findings for blood, spleen, and liver from deermice and mice and the findings with cultivated dermal fibroblasts of those species in experiments in which the time frames for the exposures to the agonists were similar. Whether this would also hold true for shorter or longer periods of exposure remains to be determined.

One of these correspondences was Slpi, which encodes an inhibitor of proteases of neutrophils and other phagocytic cells. Among its functions, the secreted protein has a role in wound repair (58) and has an inhibitory effect on the formation of neutrophil extracellular traps (59). Of relevance for a study of an infection-tolerant reservoir for the Lyme disease agent was the report that Slpi-deficient mice infected with *B. burgdorferi* had more severe arthritis and greater bacterial burdens in their joints than their wild-type counterparts (60). In prior studies we observed that expression of Slpi increased orders of magnitude in the blood of *P. leucopus* treated with LPS or infected with a relapsing fever *Borrelia* species (19). For LPS-treated mice there was only a modest increase in Slpi transcription from a low level. These observations notwithstanding, it was not a given that dermal fibroblasts of *P. leucopus* would transcribe Slpi and produce the protein, but that was the finding here. Transcription was high at baseline in the deermouse cultures and increased further with TLR2 agonist exposures (Figure 4). In contrast, with the mouse cells there was the same comparatively low expression of Slpi as observed in the in vivo experiment. While there remain questions about regulation of Slpi expression, the tight correlation of Slpi transcription across species with that of the stress responsive transcription factor Nfe2l2, more commonly known as Nrf2, indicates a direct regulatory role, as has been reported for the mouse (61). Nfe2l2 regulates the transcription of a variety cytoprotective proteins, including antioxidants, such as heme oxygenase 1 and superoxide dismutase 2, and chaperones (62). Figure 4 reveals a high constitutive level of transcription of Nfe2l2 by the *P. leucopus* fibroblasts compared to the control *M. musculus* cells. This is a characteristic of the long-lived naked mole rat, an established experimental model for aging research (63), and the comparatively long-lived Snell dwarf mutant mouse (64). Constitutive expression of Nfe2l2 has also been associated with longevity in the nematode *Caenorhabditis elegans* (65). A possible explanation of the successful serial propagation of the deermouse cells, but not the mouse cells, in presence atmospheric concentrations of oxygen was a greater capacity to withstand oxidative stress.

Another distinguishing feature of *P. leucopus* cells of the blood and spleen was the near absence of transcription of Nos2 (19, 20), which encodes the mitochondrion-based, inducible nitric oxide synthase. This when Arg1 is transcribed at high levels at baseline and even higher with LPS exposure. As noted previously for *P. leucopus* fibroblasts (19) and in the present study (Figure 3), Nos2 is transcribed constitutively and at higher levels after exposure to an agonist of TLR4 or TLR2. But this was starting at a level many-fold lower than for the *M. musculus* fibroblasts at baseline. Arg1 expression was hardly detectable in the mouse fibroblasts and substantially higher in deermouse cells under both control and treated conditions, similarly to what was observed in blood and spleen. What the in vitro study added to the picture of arginine metabolism in the two species was the differential expression of two other arginine metabolism genes associated with the mitochondrion, Arg2 and Ass1, and an arginine transporter, Slc7a2 (42), which, unlike Arg2 and Ass1, was transcribed at comparatively high levels both in controls and agonist-treated deermouse cells (Figure 3). These findings draw attention to the mitochondria and the dichotomy between *P. leucopus* and *M. musculus* in the sites of activity for arginine metabolism genes of recognized importance in immunity (66, 67).

As in the study of whole blood of *P. leucopus* and *M. musculus*, there were differences between species in transcription of ERV and TE sequences in cultured fibroblasts. In the study of LPS-treated animals this was restricted to selected retroviral envelope protein sequences, which decreased in transcription in the deermice while increasing in the mice (20). In the present study much larger sets of the ERV sequences of small to large size were used as references. These datasets were comparable between species in numbers and in the overall distributions of lengths and over the wide ranges of levels of transcription. Given the high redundancy of ERVs of all sizes in the mammalian genomes, confidently ascribing a location for the source of a ∼150 nt cDNA read is a challenge, and this limitation, which applies to all studies of this sort, is acknowledged. That is why the emphasis was on the lengths of the ERVs as a class and not their precise identification. With that disclaimer, we found that the more highly expressed ERV sequences at baseline were generally longer in *M. musculus* than in *P. leucopus* (Figure S6) and that this was also true for those sequences that were upregulated DEGs for one species or another (Figure 7).

Deermice have recently experienced massive invasions of diverse ERVs, resulting in a large expansion of KRAB zinc finger proteins (KRAB-ZFPs) responsible for ERV suppression (27, 68). This expansion of suppression machinery could enable deer mice to control ERV expression more effectively, with important implications for inflammation, immune response, and longevity (69-71). Consistent with this, previous work has shown evidence that *P. leucopus* may control ERV expression more effectively compared to other rodents (20). Our data here also generally support this interpretation. However, we also observe expression of full-length ERVs in *P. leucopus* encoding all or most of the proteins of a leukemia-type retrovirus. Interestingly, this particular ERV recently arose in the deermouse germline via interspecies horizontal gene transfer, exists at low copy numbers, and encodes a full-length env genes that is expressed, suggesting that it might still function as an infectious virus (27). Such ERVs often evade host suppression machinery so this ERV’s expression might not be too surprising. Nonetheless, these observations lead to important questions about the dynamics of ERV suppression in deermice, and how more effective ERV suppression might impact immune response to other pathogens.

A more manageable number of ERV sequences to consider were the upregulated DEGs unique to transformed *P. leucopus* cells. Among the P47 DEGs were three ERVs of over 9 kb and that would encode Env proteins as well as Gag-pol polyproteins. Activation of RNA tumor viruses in spontaneously transformed mouse cell lines with or without inciting factors has long been noted (31, 72). *P. leucopus* provides another example of this, but this phenomenon in this species remains incompletely characterized. A future step would be documentation of expression of viral proteins by these longer ERVs, such as with specific antibodies, mass spectrometry, or assay for reverse transcriptase activity. While the P47 fibroblasts behaved as if immortal, including increased expression of the oncogene Myc (Table 2), other distinguishing features of transformed cells, such as their karyotypes, have not been examined.

In distinction to these parallels between the vitro and in vivo results, an in vitro finding lacking a counterpart in the animal studies was a difference between species in transcription of the cytokine interleukin-11. In the blood, spleen, or liver of mice and deermice there was scant to no transcription of Il11 under either baseline or agonist exposure conditions (19, 20). Does the low transcription of Il11 in primary cultures of deermouse fibroblasts fairly represent what occurs in other tissues or other pathologic conditions? Re-examination of results of RNA-seq of the skin of *P. leucopus* with or without infection with *B. burgdorferi* found no detectable transcription of Il11 (6). Whether there is an association of IL-11 with aging in *Peromyscus*, as has been reported for other species (46, 47), remains to be determined.

What we can consider here is the possible source of the Il11 transcription in the fibroblasts. This was evident for the mouse fibroblasts: the right-sided cluster 12 (Figure 7). Leukemia inhibitory factor (Lif), another member of the IL-6-type cytokine family, and interleukin 1 receptor antagonist (Il1rn) also were associated with cluster 12 (Table S6). Il11, Il1rn, and Lif were more highly transcribed in P47 than in P4 deermouse cells (Table 2 and Table S7) and, unlike some chemokines, like Ccl2 and Cxcl10, unchanged in transcription in P47 cells exposed to Pam3CSK4 or LPS (Table S8). This suggests that with spontaneous transformation and possible de-differentiation there was release of repression of these genes, allowing for constitutive expression. The pro-inflammatory cytokine IL-6 also had right-side associations, namely cluster 4 of *P. leucopus* (Figure 7). But, unlike the other IL-6-type family members, Il6 transcription was much decreased in the transformed cells (Table 2), an indication of a different form of regulation.

As the distinguishing features between *P. leucopus* and *M. musculus* mount and the list lengthens, a relationship between the infection-tolerant phenotype and the greater longevity of this deermouse begins to emerge. Similar differences between deermouse and house mouse had also been noted in comparative studies with different experimental designs for either the animals themselves (73) or bone marrow-derived macrophages (74). We knew before of the low susceptibility of *P. leucopus* to what would be lethal doses of LPS for a mouse, the moderated expression of interferons and downstream genes, and a polarization profile of alternatively activated macrophages tilting toward an anti-inflammatory outcome. The present study adds these distinctions: divergent natural histories of cultured fibroblasts, constitutive expression of the stressor-response transcription factor Nfe2l2/Nrf2, diminished expression of the aging-associated cytokine IL-11, and an indisposition to transcribe long ERVs with their potential to elicit recognition by PRRs and the inflammatory consequences of that.

Can we identify ramifications of the work of particular relevance to Lyme disease, as manifested either in an experimental animal model or human illnesses as we encounter them? After all, the agonist chosen for the study is one of the identified PAMPs of *B. burgdorferi* and other Lyme disease agents of the genus. Of particular interest would be findings from the present study that might point to new lines of investigations for Lyme disease research. We think one of these is the phenomenon of widespread transcription of sequences of ERVs and TEs, both under baseline conditions and differentially after exposure to the lipopeptide PAMP in mouse and deermouse cells. The conjecture is not that there is production of whole virions; there is no evidence of that in either our studies or elsewhere in the literature. Rather, it is that translated products, such as Env protein fragments or parts of reverse transcriptases, of retrovirus-derived sequences serve as PAMPs for cell autonomous immune responses that would not be expected for infections by bacteria that are largely extracellular in the host (20). Two other topics that in our view merit further study in experimental models and in clinical research studies of Lyme disease are immunity-associated genes of arginine metabolism (66, 67) and mitigations against oxidative stress, especially by genes under the influence of the transcription factor Nrf2/Nfe2l2 (62).

## Materials and Methods

### Animals

The study was carried out in accordance with the recommendations in the National Institutes of Health’s *Guide for the Care and Use of Laboratory Animals*: Eighth Edition of the National Academy of Sciences, and according to ARRIVE Guidelines (https://arriveguidelines.org). University of California Irvine (UCI) protocols AUP-23-127 and AUP-24-042 were approved by UCI’s Institutional Animal Care and Use Committee.

*Peromyscus leucopus*, here also referred to as “deermice”, were of the outbred LL stock that originated with 38 animals captured near Linville, NC, and thereafter comprised a closed colony without sib-sib matings at the Peromyscus Genetic Stock Center at the University of South Carolina (75). Deermice used in this study were each from different mating pairs and bred at the UCI vivarium. Outbred *Mus musculus* strain CD-1 and inbred *M. musculus* FVB/NJ strain, here also referred to as “mice”, were obtained from Charles River Laboratories and Jackson Laboratory, respectively. Rodents were maintained in the AAALAC-accredited UCI vivarium, with 2 to 5 animals per cage, a 16-h light/8-h dark schedule, room temperatures of 21-23°C, and humidity of 30-70%. Food and water were available ad libitum; the diet was 2020X Soy Protein-Free Extruded Rodent Diets (Teklad). The rodents were euthanized by carbon dioxide overdose followed by exsanguination by open thorax cardiac puncture.

### Tissue culture medium

The medium was RPMI 1640 medium (Gibco) supplemented with 10% heat-inactivated fetal bovine serum (Gibco), 2 mM L-glutamine, 100 µM L-asparagine, 50 µM 2-mercaptoethanol, 250 ng/ml amphotericin B (Gibco, Fisher Scientific), 500 U/ml penicillin G, and 500 µg/ml streptomycin (76).

### Processing of tissue

Immediately after euthanasia, ∼1 cm diameter of full-thickness ear tissue was excised with sterile instruments, immersed in 70% ethanol for 5 min, and allowed to dry in a biosafety cabinet. Once dry, the tissue transferred to a 1.8 ml polystyrene screwcap tube containing 1.5 ml medium with 2 mg/ml collagenase type I (Millipore Sigma). Ear tissue was minced while in the tube with scissors down to ∼3 mm size, and then the suspension was incubated at 37°C on a horizontal shaker at 200 rpm for 75 min. The suspension was then pushed through a 70 µm cell strainer with the rubber end of a 5 ml syringe plunger into 10 ml of medium. The pass-through liquid was centrifuged at 400 x g at room temperature for 5 min. The supernatant was removed, the pellet was suspended in 10 ml medium, the suspension was centrifuged again, and the supernatant was removed.

### Tissue cultivation

After the final centrifugation, the cell pellet was suspended in 10 ml of pre-warmed medium. This was transferred to a 10 cm diameter polystyrene culture dish (Fisher Scientific FB0875712) with lid and incubated at 37°C in a humidified atmosphere with 5% CO_2_ (passage 0). After 24 h the medium was removed by aspiration, adherent cells were washed with phosphate-buffered saline, pH 7.4 (PBS) once, and then fresh medium was added at that point and then every 3-4 d as needed. The cultures were monitored by phase microscopy at 100x magnification with Olympus CK2 inverted phase contrast microscope. For sub-cultures (passage 1 onwards) when the adherent cells were 80-90% confluent, medium was aspirated, the cells were washed once with PBS. To release the cells, 2 ml of a solution of 0.02 mM trypsin and 0.48 mM sodium EDTA (trypsin-EDTA) was added before a 5 min incubation. The resultant suspension of cells were transferred to a 15 ml polystyrene conical tube containing 8 ml of medium and centrifuged at 400 x g for 5 minutes. The supernatant was removed, and the cell pellet was resuspended in 10 ml of supplemented medium. A 1:1 ratio of 0.4% w/v trypan blue in PBS and cell suspension was used for manual counting of cells with a Petroff-Hausser counting chamber under the inverted microscope at 200x magnification. Once counted, 100,000 cells were seeded into a new 10 cm petri dish with 10 ml of pre-warmed medium for an initial density of 10,000 cells/ml at time 0. The remaining cell suspension was centrifuged at 400 x g for 5 min. The cell pellet was resuspended in 1 ml of medium with 10% cell culture grade dimethyl sulfoxide and then transferred to 2 ml screwcap cryogenic vials (Greiner) and stored in liquid nitrogen. For determination of population doubling time the cultures were monitored daily until adherent cells were 80-90% confluent. After trypsinization, the harvested cells were suspended in 10 ml of medium and counts of cells per ml were made at time X, where X is the interval in days from time 0. The log_10_ of the densities at times 0 and x were plotted against time for a log-linear regression.

### Photomicrographs

Digital images were taken on an Olympus CK2 inverted phase contrast microscope equipped with a LabCam Ultra (iDu Optics) adapter for an iPhone 14 Pro (Apple Computer) and with the 15x magnification eyepiece, 10x magnification objective lens, and 2x digital zoom. The final magnification was 150x for the 4032 × 3024 resolution High Efficiency Image File Format (HEIF) format files.

### Chemicals

The lipopeptide Pam3CysSerLys4 (Pam3CSK4; Invivogen) and ion-exchange chromatography-purified lipopolysaccharide (LPS) of *Escherichia coli* O111:B4 (Sigma-Aldrich) were dissolved as stock solutions at 1 mg/ml concentration in endotoxin-free, sterile 0.9% (w/v) NaCl (saline; Sigma-Aldrich). Stock solutions were stored at -20°C in a non-frost-free freezer until the day of use, and when dilutions were freshly made in saline.

### Cell culture treatments

Cell cultures at the point of 80-90% confluency were treated with trypsin-EDTA, counted using a hemocytometer, and split into 3 wells each in a 6-well polystyrene tissue culture plate (Fisher Scientific) at 3 × 10^5^ cells per well containing 2 ml of medium. For the study of single cells there were 5 × 10^5^ cells per well. Cells were incubated at 37°C in 5% CO_2_ for 24 h prior to treatment before either saline alone or Pam3CSK4 or LPS in saline was added for final concentrations of 1 µg/ml or 10 µg/ml for Pam3CSK4 or 1 µg/ml for LPS. The suspensions were incubated at 37°C in 5% CO_2_ for 4 h, which is the interval in the previous in vivo experiments between injection of TLR agonist and termination (19, 20).

### RNA extraction

After incubation with TLR agonist or medium alone, medium was aspirated, and 300 µl of DNA/RNA Shield (Zymo Research) was added to adherent cells followed by addition of 300 µl of Zymo RNA Lysis Buffer (Zymo Research). The Quick-RNA Miniprep Plus Kit (Zymo Research) was used for the isolation of skin fibroblast RNA following manufacturer’s instructions, including for DNase treatment. RNA was eluted from the spin-column in 50 µl of Zymo Nuclease Free Water. Quantification, purity, and RNA integrity were assessed using High Sensitivity Qubit 2.0 fluorometer, Nanodrop ND-1000 spectrophotometer, and Agilent Bioanalyzer 2100. RNA integrity number (RIN) values for samples were ≥9.0.

### Bulk RNA-seq

cDNA libraries were produced with the Illumina TruSeq Stranded mRNA kit. Multiplexed libraries were sequenced at UCI’s Genomics Research and Technology Hub on an Illumina NovaSeq 6000 with paired-end chemistry, 150 cycles, and ∼120-250 million reads per sample. Read quality was analyzed by FastQC (https://www.bioinformatics.babraham.ac.uk/projects/fastqc/). Fastq files of reads were trimmed of low-quality reads (Phred score of <15), adapter sequences, and homopolymeric 5’ or 3’ ends using Trimmomatic (77). Trimmed reads were aligned to reference sets with a length fraction of 0.35, a similarity fraction of 0.9, and a cost of 3 (out of 3 maximum) each for a mismatch, insertion, or deletion to the reference sets using CLC Genomics Workbench v. 25 (Qiagen Aarhus A/S). Library size normalization was done by the TMM (trimmed mean of M values) method (78).

For the 22,654 *P. leucopus* protein coding sequences (CDS) manual annotation of 1141 gene loci without assigned gene names in the GCF_004664715.2 assembly and annotation (out of a total of 7148 with “LOCxxxxxxxxx” designations) had been carried out on ad hoc basis in those cases when a locus was identified as transcribed and a candidate DEG by RNA-seq in the present or a previous study (79). For the 22,760 protein coding genes identified in the *M. musculus* C57BL/6 reference genome transcripts (GCF_000001635.27_GRCm39), there were 97,760 isoforms. For 3927 loci there was a single listed isoform, 13,786 loci with two isoforms, and 6047 with three or more. For comparability with the *P. leucopus* dataset, the first listed isoform in the *M. musculus* reference genome CDS dataset was used (80). To assess whether other isoforms besides the first listed for *M. musculus* would provide additional or different results by RNA-seq, the 13,786 CDS with two listed isoforms were used. The sequences for these were extracted as separate sets of isoform 1 and isoform 2 for each of these CDS (Dryad Table D7) and subsequently used for independent RNA-seq and differential gene expression analysis.

For fold-change (FC) analyses of results that paired by individual *P. leucopus* (n = 22,654 CDS) or *M. musculus* (n = 22,760 CDS) cell lines, unique counts were first normalized by total reads for the corresponding sample across the species and then adjusted as reads per kilobase (Dryad Tables D1 and D2). Paired *t*-tests were performed with pairing of the same cell line under the different conditions. For cross-species comparisons of the 14,979 CDS in common and with assigned gene names (Dryad Table D3) the ratio of length-adjusted, total count-normalized reads for a given gene to the house keeping gene Gapdh was used. The mean (standard deviation) log_10_ Gapdh normalized unique counts were 4.8 (0.05) across 15 *P. leucopus* samples (i.e. 5 cell lines x 3 conditions) and 5.1 (0.05) across 15 *M musculus* samples. There was little or no discernible effect of the treatments on transcription of Gapdh: the mean (standard deviation) paired fold-changes (paired *t*-test *p* value) of Pam3CSK4 treated cells to control were 1.08 (0.16), 1.12 (0.08), 1.00 (0.90), and 1.13 (0.07) for mouse 1 µg/ml, mouse 10 µg/ml, deermouse 1 µg/ml, and deermouse 10 µg/ml sets, respectively (Dryad Table D3). Following the recommendation of Hedges et al. we used the natural logarithm (LN) of ratios (81).

For the input data for gene ontology term analysis (see below) differential gene expression (DEG) within a species and across conditions (i.e. without pairing by cell line) transcripts per million (TPM) were used as the expression level. The DEG analysis was with the CLC Genomics Workbench suite’s Differential Expression Analysis tool, which is similar to EdgeR and DESeq2 in assuming a negative binomial distribution for expression levels and fits a separate Generalized Linear Model for each (82). There is adjustment for dispersion (76), and for this analysis it was without downweighting of outliers.

### Gene ontology (GO) term analysis

The analysis was implemented with the tools of Metascape (https://metascape.org) (83) with default settings and *Mus musculus* as the closest taxon for comparison. Similarity matrices were hierarchically clustered, and a 0.3 similarity threshold was applied to trim resultant trees into separate clusters. The lowest *p* value term represented each cluster shown in the heatmaps. Besides the terms beginning with ‘GO’ of the Gene Ontology resource (http://geneontology.org) (84), other terms refer to Kegg Pathway database (https://www.kegg.jp) for ‘mmu.’ designations, WikiPathways database (https://www.wikipathways.org) for ‘WP…’ designations, and Reactome database (https://reactome.org) for ‘R-MMU…’ designations.

### Transposable element sequences

We annotated endogenous retrovirus (ERV) and other transposable elements (TE) in both the *Mus musculus* C57BL/6 and *Peromyscus leucopus* genomes using a combined nonredundant database of lineage-specific and ancestral TEs in each species. To do this, we first retrieved curated TE models for *P. maniculatus* and *Mus musculus* from DFAM (https://doi.org/10.1186/s13100-020-00230-y), as well as all ancestral TE models for each species. Then, we clustered redundant models using CD-HIT-EST (version 4.8.1) (https://doi.org/10.1093/bioinformatics/bts565) with these parameters: -n 10 -c .8 -r 1 -i. To annotate TEs in each genome using our combined TE library as input, we employed RepeatMasker (version 4.1.2) (https://www.repeatmasker.org) with these parameters: -pa 12 -excln -s -no_is -u -noisy -html -xm -a -xsmall. The names, genome locations, and lengths for identified ERVs and TEs are provided in Dryad Table D5 for *M. musculus* and Table D6 for *P. leucopus*. For bulk RNA-seq analysis with these sequences as the reference sets, the measure of expression was TPM instead of unique reads. We wrote a custom python script (Supplementary_Text_3) to identify in these sub-sets of ERV sequences those ORFs that had an ATG as start codon (where A is position +1), were at least 30 codons, and had both a purine at position -3 and a G at position +4. Identified protein products of the ORFs meeting these criteria were used for BLASTP searches (https://blast.ncbi.nlm.nih.gov/Blast.cgi) with default settings of non-redundant protein sequences for *Mus musculus* and gammaretroviruses of the GenBank database (https://www.ncbi.nlm.nih.gov/genbank/).

### Single-cell RNA-seq

After incubation of 5 × 10^5^ second passage *P. leucopus* fibroblast cells or second passage *M. musculus* strain FVB/NJ fibroblast cells with medium and saline alone, medium with 10 µg/ml Pam3CSK4, or medium with 1 µg/ml LPS, the medium was aspirated from the wells. The source animals were female and ∼220 days of age. Adherent cells were washed with 2 ml of PBS. A 0.5 ml volume of trypsin-EDTA solution was added to each well, and the cells were incubated at 37°C in 5% CO_2_ for 5 min. Detached cells from each condition were pooled into 15 ml polystyrene conical tubes by species, and 10 ml of medium was added to neutralize trypsin activity. The pooled cell suspensions for *P. leucopus* and *M. musculus* were centrifuged at 400 x g for 5 minutes and supernatant was removed. Cells of pellets were resuspended in Illumina Cell Suspension Buffer of the Single Cell 3’ RNA Prep Kit (Illumina). Cell viability and counts were determined in triplicate using trypan blue stain and a hemocytometer counting chamber. Cells were diluted in Cell Suspension Buffer to a concentration of 5000 per µl concentration and a final total count was determined. Approximately 40,000 cells for each species were further processed using an Illumina Single Cell 3’ RNA Prep Kit for mRNA capture, cDNA synthesis, and library preparation, according to manufacturer’s instructions. cDNA quality control and fragment analysis were performed using a DNA High Sensitivity chip on an Agilent BioAnalyzer 2100. A post-library preparation quality control, fragment analysis, and KAPA library quantification (Roche) by quantitative polymerase chain reaction were performed. cDNA libraries were sequenced on an Illumina NovaSeq X Plus system with paired-end reads, 150 cycles, and incorporation of a 2% PhiX spike-in. The yields were 1.6 × 10^9^ and 1.8 × 10^9^ reads for *M. musculus* and *P. leucopus*, respectively.

### Single-cell RNA-seq analysis

The raw fastq files were processed using DRAGEN v 4.4.2 software on the BaseSpace Sequencing Hub (Illumina). The reference genomes used were default GCF_000001635.20 genome on the site for *M. musculus* and an imported genome (GCF_004664715.2) for *P. leucopus*. At the data analysis stage for the *P. leucopus* reference set, updated annotations with formal gene names (80) substituted for loci with incomplete descriptions in the original annotation. The DRAGEN output included filtered matrix, features, and barcodes files, which were subsequently analyzed in *R* (85) using Seurat v5.2.1 (https://satijalab.org/seurat) (86). Seurat facilitated Uniform Manifold Approximation and Projection (UMAP) clustering via principal component analysis (PCA), visualization of gene expression on UMAPs through feature plots, cluster modifications, and identification of the enriched genes for each cluster. For the UMAP, principal components were used with the Seurat FindNeighbors setting from 1:10 and the FindClusters resolution set to 0.5. For the single-cell RNA-seq data the criteria for GO term analysis of signifying genes of clusters were an enrichment of ≥1.5 and *p* value <10^−4^ (Table S6).

### MALDI-TOF mass spectrometry

After aspiration of medium, adherent *P. leucopus* dermal fibroblasts of low or high passage grown on 10 cm diameter polystyrene petri dishes were washed with 5 ml PBS. Adherent cells were lysed by the addition of 1 ml of Radioimmunoprecipitation Assay buffer with protease inhibitor (ThermoFisher). The petri dish bottom was scraped with a cell scraper and homogenized. Cell lysate suspension was transferred to a sterile 1.5 ml polystyrene screwcap tube, and sonication was performed with alternating cycles of 15 s of sonication and 15 s on ice over 1 min. Lysates were then centrifuged at 21,000 x g for 10 min at room temperature, and the supernatant was decanted. The lysate was subjected in sodium dodecyl sulfate polyacrylamide gel electrophoresis on a 4-12% gradient Bis-Tris denaturing polyacrylamide gel on a NuPAGE apparatus (Invitrogen). electrophoresis was performed, and Coomassie blue-stained bands of the desired size range were excised. In-gel digestion with trypsin was carried out as described by Shevchenko et al. (87). In brief, destaining was performed with a 1:1 acetonitrile and ammonium bicarbonate, followed by reduction with dithiothreitol, and alkalization with iodoacetamide. An overnight trypsin digestion at 37°C was carried out, and this was followed by extraction of the mixture of peptides, spotting them onto a plate reader, and air drying. Cell analytes were measured using matrix-assisted laser desorption/ionization (MALDI) on a Bruker UltrafleXtreme instrument at the UCI Department of Chemistry Mass Spectrometry Facility. The ribosomal protein S18 (Rps18; XP_028744221.1) and its known amino acid sequence served as a control for the targeted analysis of secretory leukocyte peptidase inhibitor (Slpi). The reference for Slpi was the processed protein (amino acids 26-131) after cleavage of the signal peptide (XP_028724460.1). Spectra were acquired using a mass range from 800-6000 Daltons. FlexControl software v. 4.0 (Bruker) was used for data collection and raw spectral data were imported into mMASS v. 5.5 (https://github.com/xxao/mMass) (88). Baseline correction and peak picking were performed using mMASS default settings.

### Statistics

Unless otherwise stated, means are presented with 95% confidence intervals in parentheses. For data that was not normally distributed, these were with asymmetrical confidence intervals. Parametric (*t* test) and non-parametric (Kruskal-Wallis) tests of significance were 2-tailed. Unless otherwise stated, the paired *t*-test *p* value is given. Categorical variables were assessed by 2-tailed Fisher’s exact test. False discovery rate (FDR) correction of *p* values for multiple testing was by the Benjamini-Hochberg method (87), as implemented with False Discovery Rate Online Calculator (https://tools.carbocation.com/FDR). Linear regression, coefficient of determination (*R*^*2*^), Multidimensional Scaling (MDS), and General Linear Model (GLM) analyses were performed with SYSTAT v. 13.1 software (Systat Software, Inc). Box plots with whiskers display the minimum, first quartile, median, third quartile, and maximum.

### Data resources

Table S1 includes links to BioProject and BioSample numbers for samples of the *P. leucopus* and *M. musculus* comparison, along with accession numbers for raw reads (SRA) and RNA-seq analysis (GEO). The low and high passage *P. leucopus* dermal fibroblast study is registered under BioProject PRJNA1169383 with BioSamples SAMN45236449-54, SRA accessions SRR31668268-73, and GEO accessions GSM9100082-87. The single cell RNA-seq study is registered under BioProject PRJNA1277016 with BioSamples SAMN49093172 for *P. leucopus* and SAMN49093173 for *M. musculus* dermal fibroblasts. The corresponding SRA accessions were SRR33982422 and SRR33982421, respectively. For this study the 5’ UTR and coding sequence for the interleukin-11 gene Il11 of *P. leucopus* LL stock was determined by sequencing of the mRNA using the RNA-seq reads. This was deposited at GenBank under accession PV639634; it corrects an error in the predicted protein coding sequence in the NCBI automated annotation for accession XM_037199381.1. Tables of large datasets deposited with the Dryad (https://datadryad.org) public repository and denoted in the text by the convention “Dryad Table Dx” are accessible at https://doi.org/10.5061/dryad.m905qfvdq. Descriptions of Dryad datasets are in Supplementary Materials (Supplementary_Text_2) and in references (79) and (80).

## Supporting information

zipped archive of supplemental figures, tables, and text

## Acknowledgements

The research was supported by National Institutes of Health grant AI-157513 (A.G.B. and A.D.L.). This work utilized resources of the UCI Genomics Research and Technology Hub (GRT Hub), parts of which are supported by NIH grants to the Comprehensive Cancer Center (P30CA-062203) and to the UCI Skin Biology Resource Based Center (P30AR075047), as well as to the GRT Hub for instrumentation (1S10OD010794-01 and 1S10OD021718-01). Some computations for this work were run on the Cannon cluster supported by the Faculty of Arts and Sciences Division of Science Research Computing Group at Harvard University.

## References

1. Hagai T, Chen X, Miragaia RJ, Rostom R, Gomes T, Kunowska N, Henriksson J, Park J-E, Proserpio V, Donati G, Bossini-Castillo L, Vieira Braga FA, Naamati G, Fletcher J, Stephenson E, Vegh P, Trynka G, Kondova I, Dennis M, Haniffa M, Nourmohammad A, Lässig M, Teichmann SA. Gene expression variability across cells and species shapes innate immunity. Nature (2018) 563(7730):197–202. doi: 10.1038/s41586-018-0657-2.

2. Zhao Y, Seluanov A, Gorbunova V. Revelations about aging and disease from unconventional vertebrate model organisms. Annu Rev Genet (2021) 55:135–59. Epub 20210820. doi: 10.1146/annurev-genet-071719-021009.

3. Bedford NL, Hoekstra HE. Peromyscus mice as a model for studying natural variation. Elife (2015) 4:e06813. Epub 2015/06/18. doi: 10.7554/eLife.06813.

4. Havighorst A, Crossland J, Kiaris H. Peromyscus as a model of human disease. Semin Cell Devel Biol(2017) 61:150–5. doi: 10.1016/j.semcdb.2016.06.020.

5. Dewey MJ, Dawson WD. Deer mice: “The drosophila of North American mammalogy”. Genesis (2001) 29(3):105–9. Epub 2001/03/17.

6. Long A, Baldwin-Brown J, Tao Y, Cook V, Balderrama-Gutierrez G, Corbett-Detig R, Mortazavi A, Barbour A. The genome of Peromyscus leucopus, natural host for lyme disease and other emerging infections. Sci Advances (2019) 5(7):eaaw6441. doi: 10.1126/sciadv.aaw6441.

7. Long PN, Cook VJ, Majumder A, Barbour AG, Long AD. The utility of a closed breeding colony of Peromyscus leucopus for dissecting complex traits. Genetics (2022) 221(1). doi: 10.1093/genetics/iyac026.

8. Barbour AG. Infection resistance and tolerance in Peromyscus spp., natural reservoirs of microbes that are virulent for humans. Semin Cell Devel Biol (2017) 61(1):115–22. Epub 2016/07/07. doi: 10.1016/j.semcdb.2016.07.002.

9. Råberg L, Graham AL, Read AF. Decomposing health: tolerance and resistance to parasites in animals. Philos Trans R Soc Lond B Biol Sci (2009) 364(1513):37–49. Epub 2008/10/18. doi: 10.1098/rstb.2008.0184.

10. Ayres JS, Schneider DS. Tolerance of infections. Ann Rev Immunol (2012) 30:271–94.

11. Medzhitov R, Schneider DS, Soares MP. Disease tolerance as a defense strategy. Science (2012) 335(6071):936–41. Epub 2012/03/01. doi: 10.1126/science.1214935.

12. Soares MP, Teixeira L, Moita LF. Disease tolerance and immunity in host protection against infection. Nature reviews Immunology (2017) 17(2):83–96. Epub 20170103. doi: 10.1038/nri.2016.136.

13. Cohn O, Yankovitz G, Peshes-Yaloz N, Steuerman Y, Frishberg A, Brandes R, Mandelboim M, Hamilton JR, Hagai T, Amit I, Netea MG, Hacohen N, Iraqi FA, Bacharach E, Gat-Viks I. Distinct gene programs underpinning disease tolerance and resistance in influenza virus infection. Cell Syst (2022) 13(12):1002-15.e9. Epub 20221213. doi: 10.1016/j.cels.2022.11.004.

14. Sacher GA, Hart RW. Longevity, aging and comparative cellular and molecular biology of the house mouse, Mus musculus, and the white-footed mouse, Peromyscus leucopus. Birth Defects Orig Artic Ser (1978) 14(1):71–96. Epub 1978/01/01.

15. Cohen BJ, Cutler RG, Roth GS. Accelerated wound repair in old deer mice (Peromyscus maniculatus) and white-footed mice (Peromyscus leucopus). J Gerontol (1987) 42(3):302–7. doi: 10.1093/geronj/42.3.302.

16. Pei G, Balkema-Buschmann A, Dorhoi A. Disease tolerance as immune defense strategy in bats: one size fits all? PLoS Pathog (2024) 20(9):e1012471. Epub 20240905. doi: 10.1371/journal.ppat.1012471.

17. Gorbunova V, Seluanov A, Kennedy BK. The world goes bats: living longer and tolerating viruses. Cell Metabol (2020) 32(1):31–43. doi: 10.1016/j.cmet.2020.06.013.

18. Downie AE, Barre RS, Robinson A, Yang J, Chen YH, Lin JD, Oyesola O, Yeung F, Cadwell K, Loke P, Graham AL. Assessing immune phenotypes using simple proxy measures: promise and limitations. Discov Immunol (2024) 3(1):kyae010. Epub 20240628. doi: 10.1093/discim/kyae010.

19. Balderrama-Gutierrez G, Milovic A, Cook Vanessa J, Islam MN, Zhang Y, Kiaris H, Belisle John T, Mortazavi A, Barbour Alan G. An infection-tolerant mammalian reservoir for several zoonotic agents broadly counters the inflammatory effects of endotoxin. mBio (2021) 12(2):10.1128/mbio.00588-21. doi: 10.1128/mbio.00588-21.

20. Milovic A, Duong JV, Barbour AG. The infection-tolerant white-footed deermouse tempers interferon responses to endotoxin in comparison to the mouse and rat. Elife (2024) 12. Epub 20240109. doi: 10.7554/eLife.90135.

21. Machtinger ET, Williams SC. Practical guide to trapping Peromyscus leucopus (rodentia: Cricetidae) and Peromyscus maniculatus for vector and vector-borne pathogen surveillance and ecology. J Insect Sci (2020) 20(6). doi: 10.1093/jisesa/ieaa028.

22. Zhong X, Lundberg M, Råberg L. Comparison of spleen transcriptomes of two wild rodent species reveals differences in the immune response against Borrelia afzelii. Ecol Evol (2020) 10(13):6421–34. Epub 20200525. doi: 10.1002/ece3.6377.

23. Sinsky RJ, Piesman J. Ear punch biopsy method for detection and isolation of Borrelia burgdorferi from rodents. J Clin Microbiol (1989) 27(8):1723–7. doi: 10.1128/jcm.27.8.1723-1727.1989.

24. Zawada SG, von Fricken ME, Weppelmann TA, Sikaroodi M, Gillevet PM. Optimization of tissue sampling for Borrelia burgdorferi in white-footed mice (Peromyscus leucopus). PLoS One (2020) 15(1):e0226798. Epub 20200124. doi: 10.1371/journal.pone.0226798.

25. Råberg L, Sim D, Read AF. Disentangling genetic variation for resistance and tolerance to infectious diseases in animals. Science (2007) 318(5851):812–4. doi: 10.1126/science.1148526.

26. Oosting M, Ter Hofstede H, Sturm P, Adema GJ, Kullberg BJ, van der Meer JW, Netea MG, Joosten LA. TLR1/TLR2 heterodimers play an important role in the recognition of Borrelia spirochetes. PLoS One (2011) 6(10):e25998. Epub 20111005. doi: 10.1371/journal.pone.0025998.

27. Gozashti L, Feschotte C, Hoekstra HE. Transposable element interactions shape the ecology of the deer mouse genome. Mol Biol Evol (2023) 40(4):msad069. doi: 10.1093/molbev/msad069.

28. Hayflick L, Plotkin SA, Norton TW, Koprowski H. Preparation of poliovirus vaccines in a human fetal diploid cell strain. Am J Hyg (1962) 75:240–58. doi: 10.1093/oxfordjournals.aje.a120247.

29. Parrinello S, Samper E, Krtolica A, Goldstein J, Melov S, Campisi J. Oxygen sensitivity severely limits the replicative lifespan of murine fibroblasts. Nat Cell Biol (2003) 5(8):741–7. doi: 10.1038/ncb1024.

30. Todaro GJ, Green H. Quantitative studies of the growth of mouse embryo cells in culture and their development into established lines. J Cell Biol (1963) 17(2):299–313. doi: 10.1083/jcb.17.2.299.

31. Aaronson SA, Hartley JW, Todaro GJ. Mouse leukemia virus: “spontaneous” release by mouse embryo cells after long-term in vitro cultivation. Proc Natl Acad Sci U S A (1969) 64(1):87–94. doi: 10.1073/pnas.64.1.87.

32. Szeto A, Sun-Suslow N, Mendez AJ, Hernandez RI, Wagner KV, McCabe PM. Regulation of the macrophage oxytocin receptor in response to inflammation. Am J Physiol Endocrinol Metab (2017) 312(3):E183–e9. Epub 20170103. doi: 10.1152/ajpendo.00346.2016.

33. Wirtz S, Becker C, Fantini MC, Nieuwenhuis EE, Tubbe I, Galle PR, Schild HJ, Birkenbach M, Blumberg RS, Neurath MF. EBV-induced gene 3 transcription is induced by TLR signaling in primary dendritic cells via NF-kappa B activation. J Immunol (2005) 174(5):2814–24. doi: 10.4049/jimmunol.174.5.2814.

34. Hagiwara K, Kikuchi T, Endo Y, Huqun, Usui K, Takahashi M, Shibata N, Kusakabe T, Xin H, Hoshi S, Miki M, Inooka N, Tokue Y, Nukiwa T. Mouse SWAM1 and SWAM2 are antibacterial proteins composed of a single whey acidic protein motif. J Immunol (2003) 170(4):1973–9. doi: 10.4049/jimmunol.170.4.1973.

35. Perez CJ, Dumas A, Vallières L, Guénet JL, Benavides F. Several classical mouse inbred strains, including DBA/2, NOD/Lt, FVB/N, and SJL/J, carry a putative loss-of-function allele of Gpr84. The Journal of heredity (2013) 104(4):565–71. Epub 20130424. doi: 10.1093/jhered/est023.

36. Foster AM, Baliwag J, Chen CS, Guzman AM, Stoll SW, Gudjonsson JE, Ward NL, Johnston A. IL-36 promotes myeloid cell infiltration, activation, and inflammatory activity in skin. J Immunol (2014) 192(12):6053–61. Epub 20140514. doi: 10.4049/jimmunol.1301481.

37. Liu C, Xu Z, Gupta D, Dziarski R. Peptidoglycan recognition proteins: a novel family of four human innate immunity pattern recognition molecules. J Biol Chem (2001) 276(37):34686–94. Epub 20010718. doi: 10.1074/jbc.M105566200.

38. Viola A, Munari F, Sánchez-Rodríguez R, Scolaro T, Castegna A. The metabolic signature of macrophage responses. Front Immunol (2019) 10:1462. doi: 10.3389/fimmu.2019.01462

39. Murray PJ. Macrophage polarization. Ann Rev Physiol (2017) 79:541–66. doi: 10.1146/annurev-physiol-022516-034339.

40. Dowling JK, Afzal R, Gearing LJ, Cervantes-Silva MP, Annett S, Davis GM, De Santi C, Assmann N, Dettmer K, Gough DJ, Bantug GR, Hamid FI, Nally FK, Duffy CP, Gorman AL, Liddicoat AM, Lavelle EC, Hess C, Oefner PJ, Finlay DK, Davey GP, Robson T, Curtis AM, Hertzog PJ, Williams BRG, McCoy CE. Mitochondrial arginase-2 is essential for IL-10 metabolic reprogramming of inflammatory macrophages. Nat Commun (2021) 12(1):1460. Epub 20210305. doi: 10.1038/s41467-021-21617-2.

41. Qualls JE, Subramanian C, Rafi W, Smith AM, Balouzian L, DeFreitas AA, Shirey KA, Reutterer B, Kernbauer E, Stockinger S, Decker T, Miyairi I, Vogel SN, Salgame P, Rock CO, Murray PJ. Sustained generation of nitric oxide and control of mycobacterial infection requires argininosuccinate synthase 1. Cell Host Microbe (2012) 12(3):313–23. doi: 10.1016/j.chom.2012.07.012.

42. Thompson RW, Pesce JT, Ramalingam T, Wilson MS, White S, Cheever AW, Ricklefs SM, Porcella SF, Li L, Ellies LG, Wynn TA. Cationic amino acid transporter-2 regulates immunity by modulating arginase activity. PLoS Pathogen (2008) 4(3):e1000023. Epub 20080314. doi: 10.1371/journal.ppat.1000023.

43. O’Rourke SA, Shanley LC, Dunne A. The Nrf2-HO-1 system and inflammaging. Front Immunol (2024) 15:1457010. Epub 20240924. doi: 10.3389/fimmu.2024.1457010.

44. Kozak M. An analysis of 5’-noncoding sequences from 699 vertebrate messenger RNAs. Nucleic Acids Res(1987) 15(20):8125–48. doi: 10.1093/nar/15.20.8125.

45. Cook SA. The Pathobiology of Interleukin 11 in Mammalian Disease is Likely Explained by its Essential Evolutionary Role for Fin Regeneration. J Cardiovasc Transl Res (2023) 16(4):755–7. Epub 20230111. doi: 10.1007/s12265-022-10351-9.

46. Cook SA, Schafer S. Hiding in plain sight: interleukin-11 emerges as a master regulator of fibrosis, tissue integrity, and stromal inflammation. Ann Rev Med (2020) 71(1):263–76. doi: 10.1146/annurev-med-041818-011649

47. Widjaja AA, Lim WW, Viswanathan S, Chothani S, Corden B, Dasan CM, Goh JWT, Lim R, Singh BK, Tan J, Pua CJ, Lim SY, Adami E, Schafer S, George BL, Sweeney M, Xie C, Tripathi M, Sims NA, Hübner N, Petretto E, Withers DJ, Ho L, Gil J, Carling D, Cook SA. Inhibition of IL-11 signalling extends mammalian healthspan and lifespan. Nature (2024) 632(8023):157–65. Epub 20240717. doi: 10.1038/s41586-024-07701-9.

48. Lendahl U, Muhl L, Betsholtz C. Identification, discrimination and heterogeneity of fibroblasts. Nat Commun (2022) 13(1):3409. Epub 20220614. doi: 10.1038/s41467-022-30633-9.

49. Sheppard CH, Kazacos KR. Susceptibility of Peromyscus leucopus and Mus musculus to infection with Baylisascaris procyonis. J Parasitol (1997) 83(6):1104–11. Epub 1997/12/24.

50. Bourgeois JS, McCarthy JE, Turk SP, Bernard Q, Clendenen LH, Wormser GP, Marcos LA, Dardick K, Telford SR, Marques AR, Hu LT. Peromyscus leucopus, Mus musculus, and humans have distinct transcriptomic responses to larval Ixodes scapularis bites. Infect Immun 93:(4):e0006525. doi: 10.1128/iai.00065-25. Epub 2025 Mar 11..

51. Kacprzyk J, Locatelli AG, Hughes GM, Huang Z, Clarke M, Gorbunova V, Sacchi C, Stewart GS, Teeling EC. Evolution of mammalian longevity: age-related increase in autophagy in bats compared to other mammals. Aging (2021) 13(6):7998–8025. Epub 20210321. doi: 10.18632/aging.202852.

52. Alcock D, Power S, Hogg B, Sacchi C, Kacprzyk J, McLoughlin S, Bertelsen MF, Fletcher NF, O’Riain A, Teeling EC. Generating bat primary and immortalised cell-lines from wing biopsies. Sci Rep (2024) 14(1):27633. Epub 20241112. doi: 10.1038/s41598-024-76790-3.

53. Seluanov A, Hine C, Bozzella M, Hall A, Sasahara TH, Ribeiro AA, Catania KC, Presgraves DC, Gorbunova V. Distinct tumor suppressor mechanisms evolve in rodent species that differ in size and lifespan. Aging Cell (2008) 7(6):813–23. Epub 20080905. doi: 10.1111/j.1474-9726.2008.00431.x.

54. Driskell RR, Watt FM. Understanding fibroblast heterogeneity in the skin. Trends Cell Biol (2015) 25(2):92–9. Epub 20141107. doi: 10.1016/j.tcb.2014.10.001.

55. Philippeos C, Telerman SB, Oulès B, Pisco AO, Shaw TJ, Elgueta R, Lombardi G, Driskell RR, Soldin M, Lynch MD, Watt FM. Spatial and single-cell transcriptional profiling identifies functionally distinct human dermal fibroblast subpopulations. J Invest Dermatol (2018) 138(4):811–25. Epub 20180131. doi: 10.1016/j.jid.2018.01.016.

56. Tyshkovskiy A, Ma S, Shindyapina AV, Tikhonov S, Lee SG, Bozaykut P, Castro JP, Seluanov A, Schork NJ, Gorbunova V, Dmitriev SE, Miller RA, Gladyshev VN. Distinct longevity mechanisms across and within speciesand their association with aging. Cell (2023) 186(13):2929-49.e20. Epub 20230603. doi: 10.1016/j.cell.2023.05.002.

57. Ma S, Upneja A, Galecki A, Tsai YM, Burant CF, Raskind S, Zhang Q, Zhang ZD, Seluanov A, Gorbunova V, Clish CB, Miller RA, Gladyshev VN. Cell culture-based profiling across mammals reveals DNA repair and metabolism as determinants of species longevity. Elife (2016) 5. Epub 20161122. doi: 10.7554/eLife.19130.

58. Ashcroft GS, Lei K, Jin W, Longenecker G, Kulkarni AB, Greenwell-Wild T, Hale-Donze H, McGrady G, Song XY, Wahl SM. Secretory leukocyte protease inhibitor mediates non-redundant functions necessary for normal wound healing. Nat Med (2000) 6(10):1147–53. Epub 2000/10/04. doi: 10.1038/80489.

59. Zabieglo K, Majewski P, Majchrzak-Gorecka M, Wlodarczyk A, Grygier B, Zegar A, Kapinska-Mrowiecka M, Naskalska A, Pyrc K, Dubin A. The inhibitory effect of secretory leukocyte protease inhibitor (SLPI) on formation of neutrophil extracellular traps. J Leukocyte Biol (2015) 98(1):99–106.

60. Yu Q, Tang X, Hart T, Homer R, Belperron AA, Bockenstedt LK, Ring A, Nakamura A, Fikrig E. Secretory leukocyte protease inhibitor influences periarticular joint inflammation in Borrelia burgdorferi-infected mice. Elife (2025) 14. Epub 20250520. doi: 10.7554/eLife.104913.

61. Schäfer M, Farwanah H, Willrodt AH, Huebner AJ, Sandhoff K, Roop D, Hohl D, Bloch W, Werner S. Nrf2 links epidermal barrier function with antioxidant defense. EMBO Mol Med (2012) 4(5):364–79. Epub 20120302. doi: 10.1002/emmm.201200219.

62. Eisenstein A, Hilliard BK, Pope SD, Zhang C, Taskar P, Waizman DA, Israni-Winger K, Tian H, Luan HH, Wang A. Activation of the transcription factor NRF2 mediates the anti-inflammatory properties of a subset of over-the-counter and prescription NSAIDs. Immunity (2022) 55(6):1082-95.e5. Epub 20220518. doi: 10.1016/j.immuni.2022.04.015.

63. Lewis KN, Wason E, Edrey YH, Kristan DM, Nevo E, Buffenstein R. Regulation of Nrf2 signaling and longevity in naturally long-lived rodents. Proc Natl Acad Sci U S A (2015) 112(12):3722–7. Epub 20150309. doi: 10.1073/pnas.1417566112.

64. Leiser SF, Miller RA. Nrf2 signaling, a mechanism for cellular stress resistance in long-lived mice. Mol Cell Biol (2010) 30(3):871–84. Epub 20091123. doi: 10.1128/mcb.01145-09.

65. Zhang P, Zhai Y, Cregg J, Ang KK, Arkin M, Kenyon C. Stress resistance screen in a human primary cell line identifies small molecules that affect aging pathways and extend Caenorhabditis elegans’ lifespan. G3 (Bethesda) (2020) 10(2):849–62. Epub 20200206. doi: 10.1534/g3.119.400618.

66. Banoth B, Cassel SL. Mitochondria in innate immune signaling. Translational Res (2018) 202:52–68. Epub 2018/08/31. doi: 10.1016/j.trsl.2018.07.014.

67. Canè S, Geiger R, Bronte V. The roles of arginases and arginine in immunity. Nature Rev Immunol (2025) 25(4):266–84. Epub 20241017. doi: 10.1038/s41577-024-01098-2.

68. Bruno M, Maisha S, Mitra A, Costello K, Watkins-Chow D, Logsdon GA, Gambogi CW, Dumont BL, Black BE, Keane TM, Ferguson-Smith AC, Dale R, Macfarlan TS. Young KRAB-zinc finger gene clusters are highly dynamic incubators of ERV-driven genetic heterogeneity in mice. bioRxiv (2025);10.1101/2025.02.26.640358:2025.02.26.640358. doi: 10.1101/2025.02.26.640358.

69. Lima-Junior DS, Krishnamurthy SR, Bouladoux N, Collins N, Han SJ, Chen EY, Constantinides MG, Link VM, Lim AI, Enamorado M, Cataisson C, Gil L, Rao I, Farley TK, Koroleva G, Attig J, Yuspa SH, Fischbach MA, Kassiotis G, Belkaid Y. Endogenous retroviruses promote homeostatic and inflammatory responses to the microbiota. Cell (2021) 184(14):3794-811.e19. Epub 20210623. doi: 10.1016/j.cell.2021.05.020.

70. Liu X, Liu Z, Wu Z, Ren J, Fan Y, Sun L, Cao G, Niu Y, Zhang B, Ji Q, Jiang X, Wang C, Wang Q, Ji Z, Li L, Esteban CR, Yan K, Li W, Cai Y, Wang S, Zheng A, Zhang YE, Tan S, Cai Y, Song M, Lu F, Tang F, Ji W, Zhou Q, Belmonte JCI, Zhang W, Qu J, Liu G-H. Resurrection of endogenous retroviruses during aging reinforces senescence. Cell (2023) 186(2):287-304.e26. doi: 10.1016/j.cell.2022.12.017.

71. Kassiotis G. The immunological conundrum of endogenous retroelements. Ann Rev Immunol (2023) 41:99–125. doi: 10.1146/annurev-immunol-101721-033341.

72. Todaro GJ. Evolution and modes of transmission of RNA tumor viruses. Parke-Davis Award lecture. Am J Pathol (1975) 81(3):590–606.

73. Gaber, AM, Mandric, I, Nitirahardjo, C, Piontkivska, H, Hillhouse, AE, Threadgill, DW, Zelikovsky, A, Rogovskyy, AS. Comparative transcriptome analysis of Peromyscus leucopus and C3H mice infected with theLyme disease pathogen. Front Cell Infect Microbiol (2023) 13: 1115350. doi: 10.3389/fcimb.2023.1115350.

74. Wells, CC, Petnicki-Ocwieja, T, Tan, S, Bunnell, SC, Telford III, SR, Hu, LT, Bourgeois, JS. Differentiating Peromyscus leucopus bone marrow-derived macrophages for characterization of responses to Borrelia burgdorferi and lipopolysaccharide. Infect Immun (2025): 93(7): e0058124. 10.1128/iai.00581-24.

75. Joyner CP, Myrick LC, Crossland JP, Dawson WD. Deer mice as laboratory animals. ILAR (1998) 39(4):322–30. Epub 2001/06/15. http://www.ncbi.nlm.nih.gov/pubmed/11406688.

76. Khan M, Gasser S. Generating primary fibroblast cultures from mouse ear and tail tissues. JoVE (2016);10.3791/53565(107). Epub 20160110. doi: 10.3791/53565.

77. Bolger AM, Lohse M, Usadel B. Trimmomatic: a flexible trimmer for Illumina sequence data. Bioinformatics (2014) 30(15):2114–20. Epub 20140401. doi: 10.1093/bioinformatics/btu170.

78. Robinson MD, Oshlack A. A scaling normalization method for differential expression analysis of RNA-seq data. Genome Biol (2010) 11(3):R25. Epub 20100302. doi: 10.1186/gb-2010-11-3-r25.

79. Milovic, A, Barbour, AG. Protein coding sequences (CDS) of the genome of a Peromyscus leucopus (2025). 10.5061/dryad.fqz612k44.

80. Milovic A, Barbour, AG. Protein coding sequences (CDS) of the genome of a Mus musculus (2025). 10.5061/dryad.hdr7sqvwr.

81. Hedges LV, Gurevitch J, Curtis PS. The meta-analysis of response ratios in experimental ecology. Ecology (1999) 80(4):1150–6.

82. McCarthy DJ, Chen Y, Smyth GK. Differential expression analysis of multifactor RNA-Seq experiments with respect to biological variation. Nucleic Acids Res (2012) 40(10):4288–97. doi: 10.1093/nar/gks042.

83. Zhou Y, Zhou B, Pache L, Chang M, Khodabakhshi AH, Tanaseichuk O, Benner C, Chanda SK. Metascape provides a biologist-oriented resource for the analysis of systems-level datasets. Nat Commun (2019) 10(1):1523. Epub 20190403. doi: 10.1038/s41467-019-09234-6.

84. Ashburner M, Ball CA, Blake JA, Botstein D, Butler H, Cherry JM, Davis AP, Dolinski K, Dwight SS, Eppig JT, Harris MA, Hill DP, Issel-Tarver L, Kasarskis A, Lewis S, Matese JC, Richardson JE, Ringwald M, Rubin GM, Sherlock G. Gene ontology: tool for the unification of biology. The Gene Ontology Consortium. Nat Genet (2000) 25(1):25–9. Epub 2000/05/10. doi: 10.1038/75556.

85. R_Core_Team. R: A Language and Environment for Statistical Computing. In: Computing RFfS, editor. 4.4.3 ed. Vienna, Austria (2025), https://www.R-project.org.

86. Hao Y, Hao S, Andersen-Nissen E, Mauck WM, 3rd, Zheng S, Butler A, Lee MJ, Wilk AJ, Darby C, Zager M, Hoffman P, Stoeckius M, Papalexi E, Mimitou EP, Jain J, Srivastava A, Stuart T, Fleming LM, Yeung B, Rogers AJ, McElrath JM, Blish CA, Gottardo R, Smibert P, Satija R. Integrated analysis of multimodal single-cell data. Cell (2021) 184(13):3573-87.e29. Epub 20210531. doi: 10.1016/j.cell.2021.04.048.

87. Shevchenko A, Tomas H, Havlis J, Olsen JV, Mann M. In-gel digestion for mass spectrometric characterization of proteins and proteomes. Nat Protoc (2006) 1(6):2856–60. doi: 10.1038/nprot.2006.468.

88. Strohalm M, Kavan D, Novák P, Volný M, Havlíček V. mMass 3: A Cross-Platform Software Environment for Precise Analysis of Mass Spectrometric Data. Anal Chem (2010) 82(11):4648–51. doi: 10.1021/ac100818g.

89. Benjamini Y, Hochberg Y. Controlling the false discovery rate: a practical and powerful approach to multiple testing. J Roy Stat Soc Series B (Methodological) (1995):289–300.

