## Supplementary material for "Dermal fibroblast cultures recapitulate differences between deermice and mice in responses to a Toll-like receptor agonist": zipped archive of supplemental figures, tables, and text: FigureS1.pdf

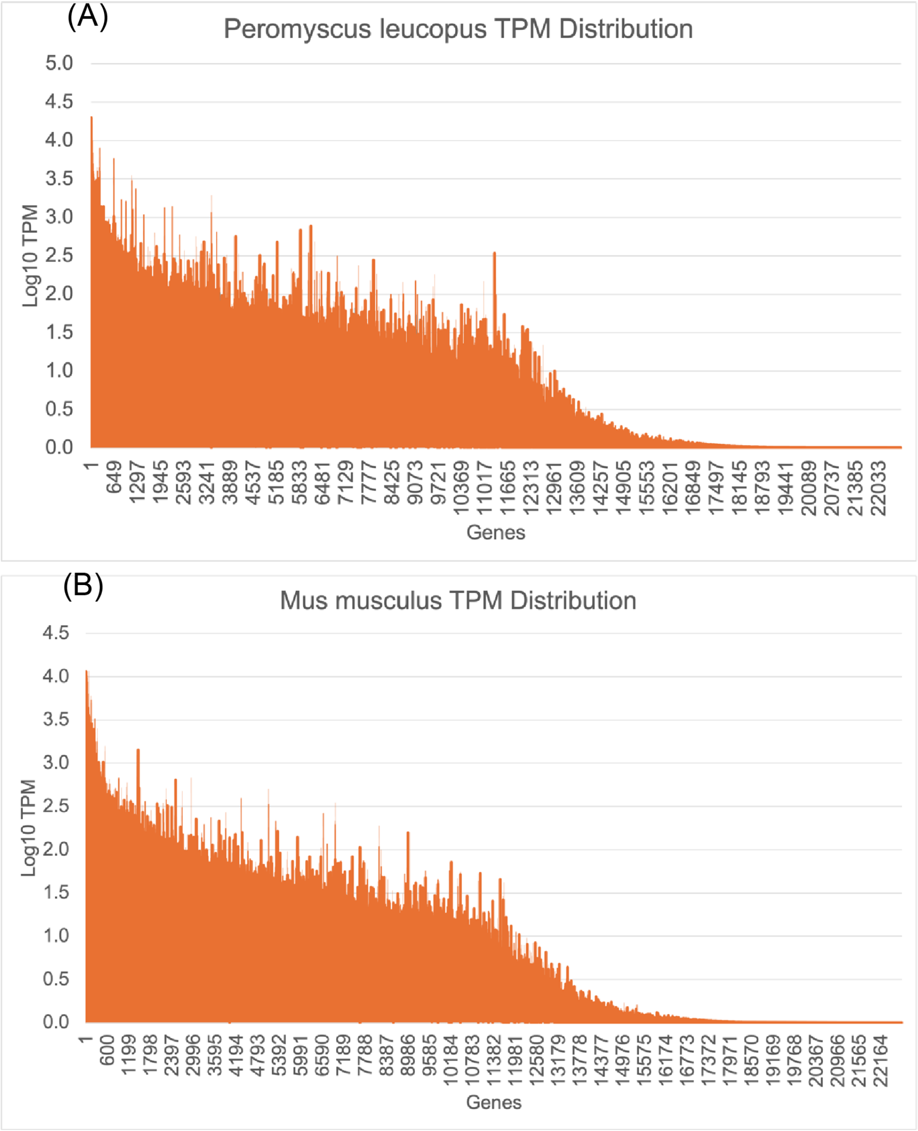


Figure S1. Histograms of the distributions of $\log_{10}$ of the mean (n=5) TPM values for 22,760 CDS of *Mus musculus* (panel A) and 22,654 CDS of *Peromyscus leucopus* (panel B) from genome-wide RNA-seq of primary dermal fibroblast cultures under control conditions. The *x*-axis is the cumulative CDS count. Data for analysis are in Dryad Tables D1 and D2.
