## Supplementary material for "Dermal fibroblast cultures recapitulate differences between deermice and mice in responses to a Toll-like receptor agonist": zipped archive of supplemental figures, tables, and text: FigureS2.pdf

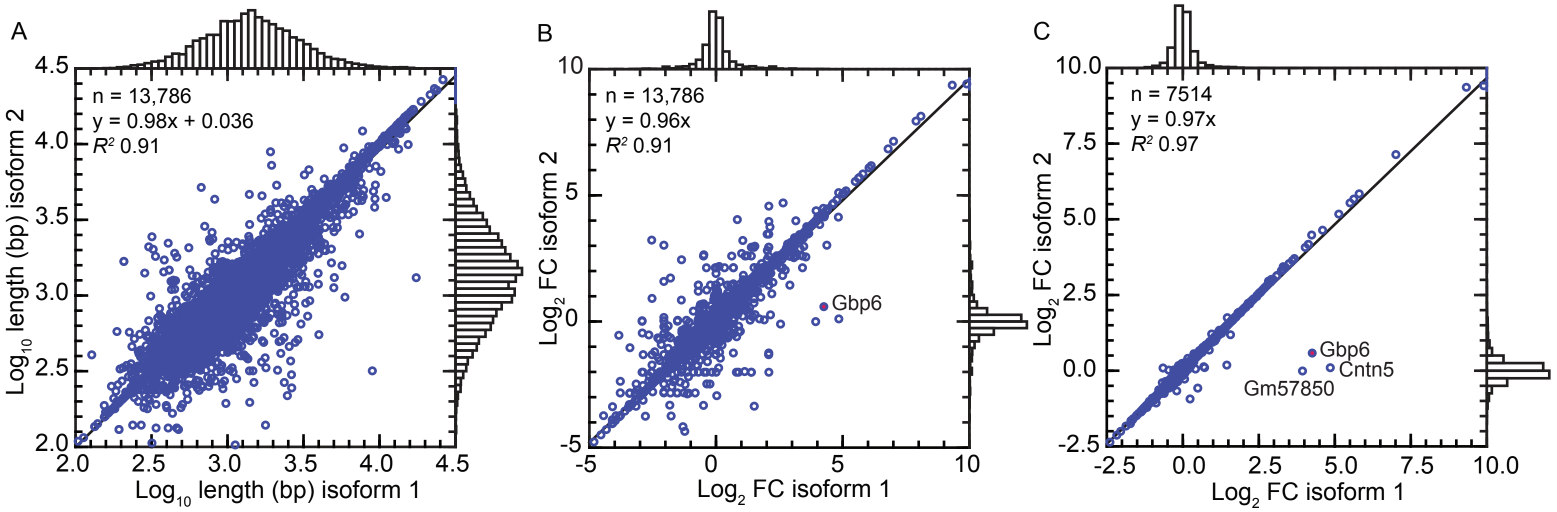


Figure S2

Figure S2. Scatter plots of bulk RNA-seq results for five Pam3CSK4-treated or untreated low passage dermal fibroblast cultures using as reference sets 13,786 *Mus musculus* protein coding sequences (CDS) for which there were two isoforms. For each panel there are provided the number of data points, univariate distributions on *x*- and *y*-axes, regression lines, equations for regression lines, and coefficients of determination ($R^2$). Panel A is a plot of log-transformed lengths in base pairs (bp) of isoform 2 on isoform 2. Panel B is a plot of log-transformed fold change (FC) of treated over untreated cells for the entire sets of sequences. Panel C is a similar plot but limited to the 7514 CDS for which the mean TPM for the treated cultures with the second isoform reference set was ≥10. In both panels B and C the coordinates for Gbp6 are indicated by label and dark red fill. Gbp6 was a DEG with isoform 1 but not isoform 2. The two other genes (Gm57850 and Cntn5) whose locations on the plots are labeled had FDR *p* values >0.05 for the first isoform set as well as the second. Data for analysis are in Dryad Table D7.
