## Supplementary material for "Dermal fibroblast cultures recapitulate differences between deermice and mice in responses to a Toll-like receptor agonist": zipped archive of supplemental figures, tables, and text: FigureS3.pdf

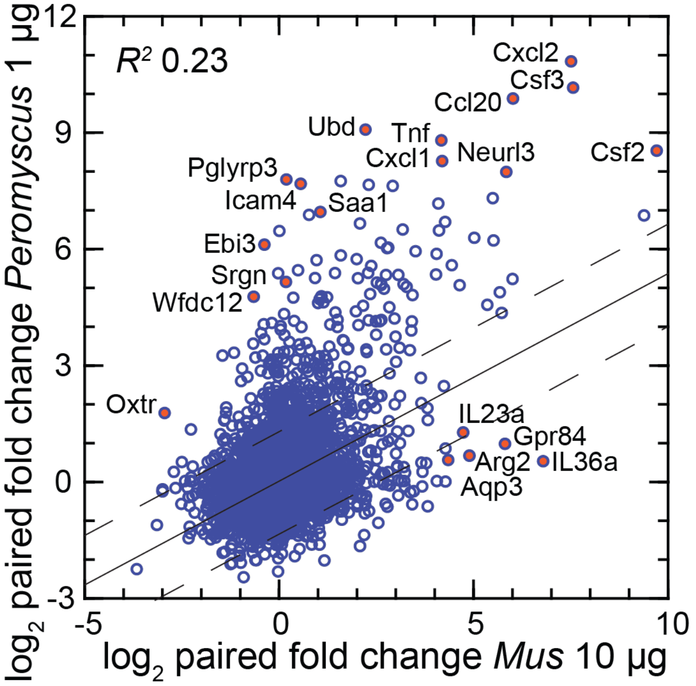


Figure S3

Figure S3. Scatter plot of log-transformed FC values for each species against each other as described for Figure 2. The difference is that the comparison for *P. leucopus* is with *M. musculus* fibroblasts exposed to the 10 µg/ml concentration of Pam3CSK4. The linear regression line with 95% confidence interval and coefficient of determination ($R^2$) are shown. Selective genes that are up-regulated and differentially expressed (DEGs) for each species are indicated by name and red fill. Data for analysis are in Dryad Table D3.
