## Supplementary material for "Dermal fibroblast cultures recapitulate differences between deermice and mice in responses to a Toll-like receptor agonist": zipped archive of supplemental figures, tables, and text: FigureS4.pdf

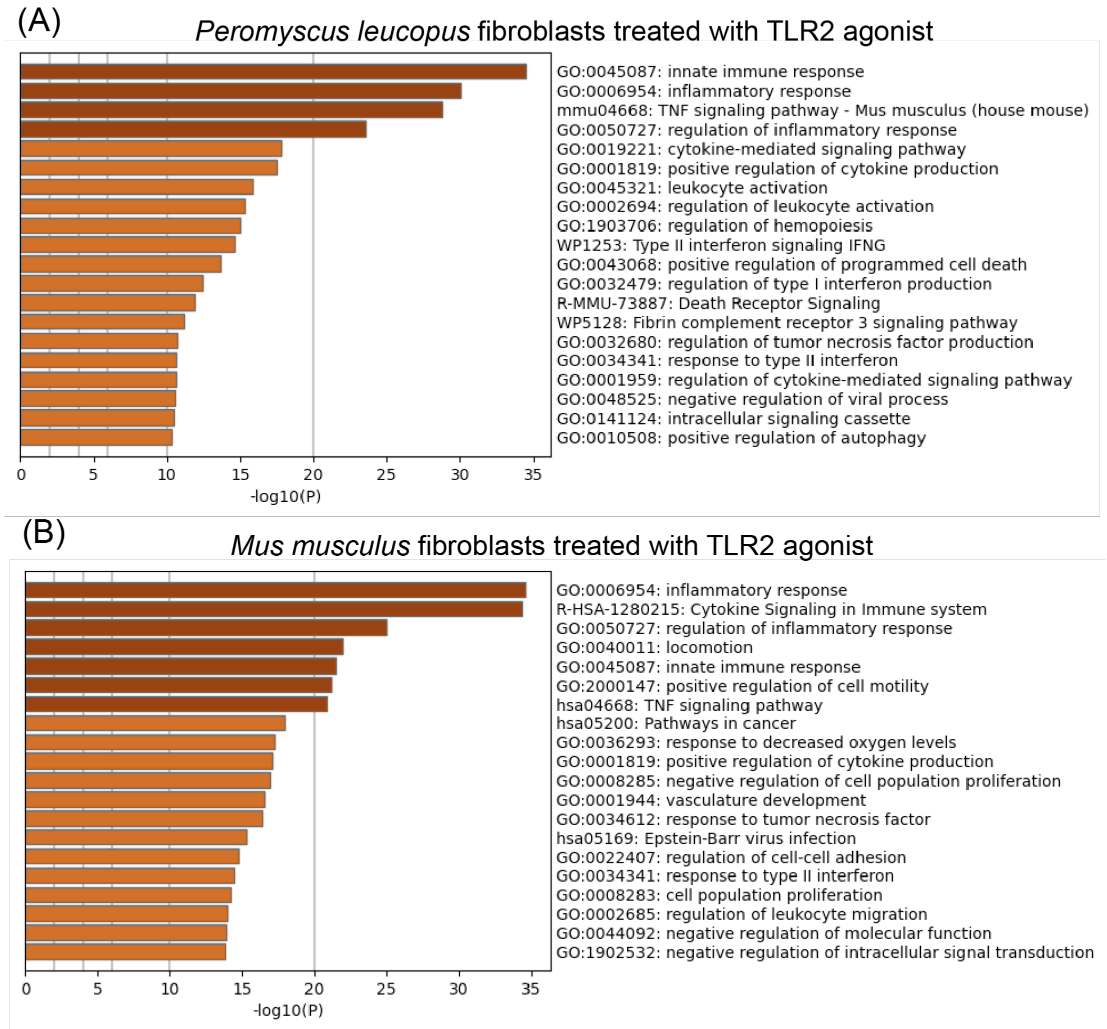


Figure S4. Horizontal bar graphs for the comparison of *Peromyscus leucopus* (panel A) and *Mus musculus (panel B)* in their responses to the TLR2 agonist Pam3CSK4 by RNA-seq and categorization of DEGs by Gene Ontology (GO) term enrichment. The analyses shown are of those sets of genes of each species that were higher in transcription in the treated cells than in the control cells. Source of data for analysis provided in Table S1 under GEO accession numbers.
