## Supplementary material for "Dermal fibroblast cultures recapitulate differences between deermice and mice in responses to a Toll-like receptor agonist": zipped archive of supplemental figures, tables, and text: FigureS5.pdf

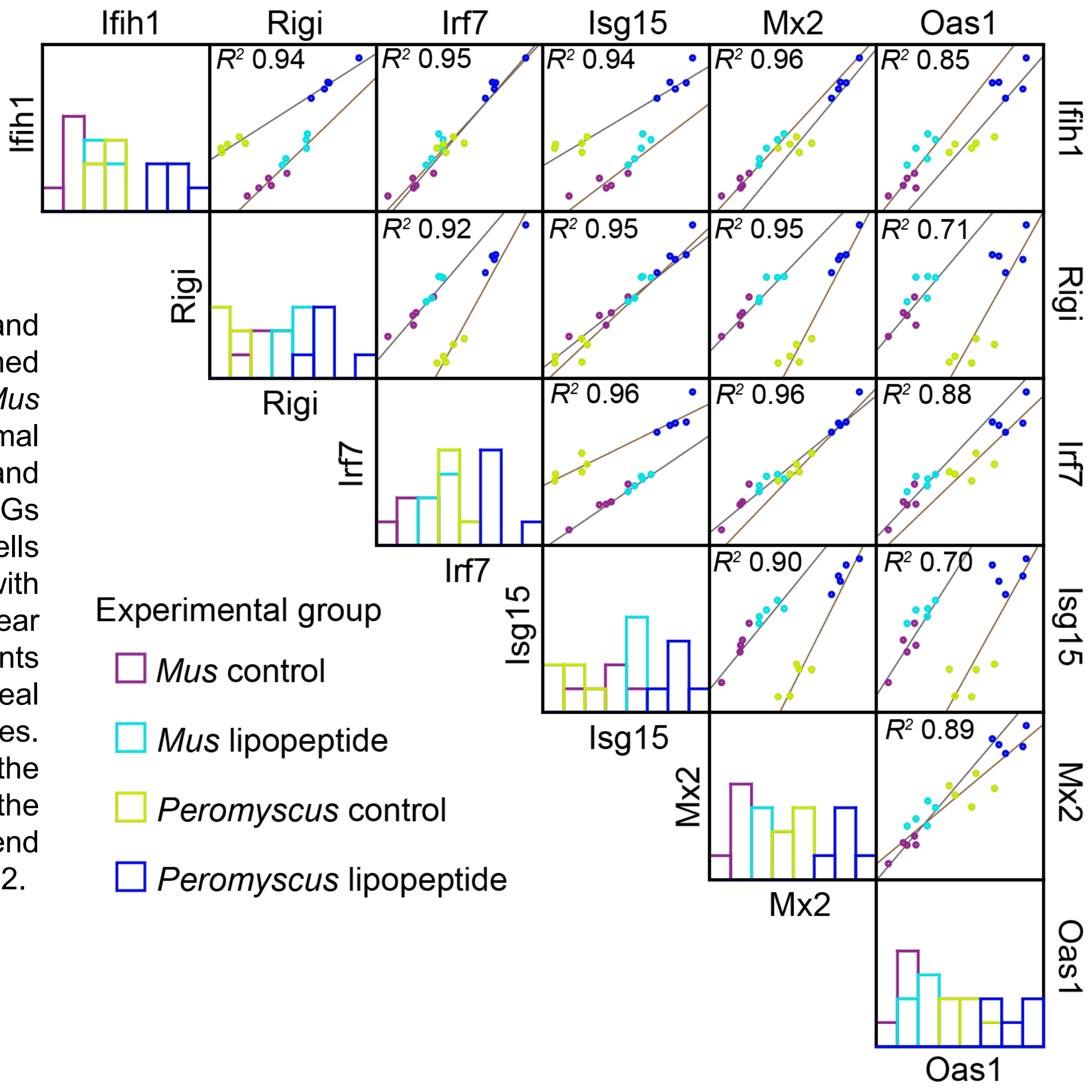


Figure S5. Matrix of scatter plots and linear regressions of log-transformed Gapdh-normalized transcriptions by *Mus musculus* or *Peromyscus leucopus* dermal fibroblasts of the PRRs Ifih1 (MDA5) and Rigi, transcription factor Irf7, and the ISGs Isg15, Mx2, and Oas1 for all pairs. The cells were without (control) or with treatment with the lipopeptide Pam3CSK4. The linear regressions are by species. The coefficients of determination ($R^2$) are for Geneal Linear Model calculations for both species. Histograms of density distributions of the values by experimental group for the variable of each *y*-axis are show at the left-end of each row. Data for analyses are in Table S2.
