## Supplementary material for "Dermal fibroblast cultures recapitulate differences between deermice and mice in responses to a Toll-like receptor agonist": zipped archive of supplemental figures, tables, and text: FigureS6.pdf

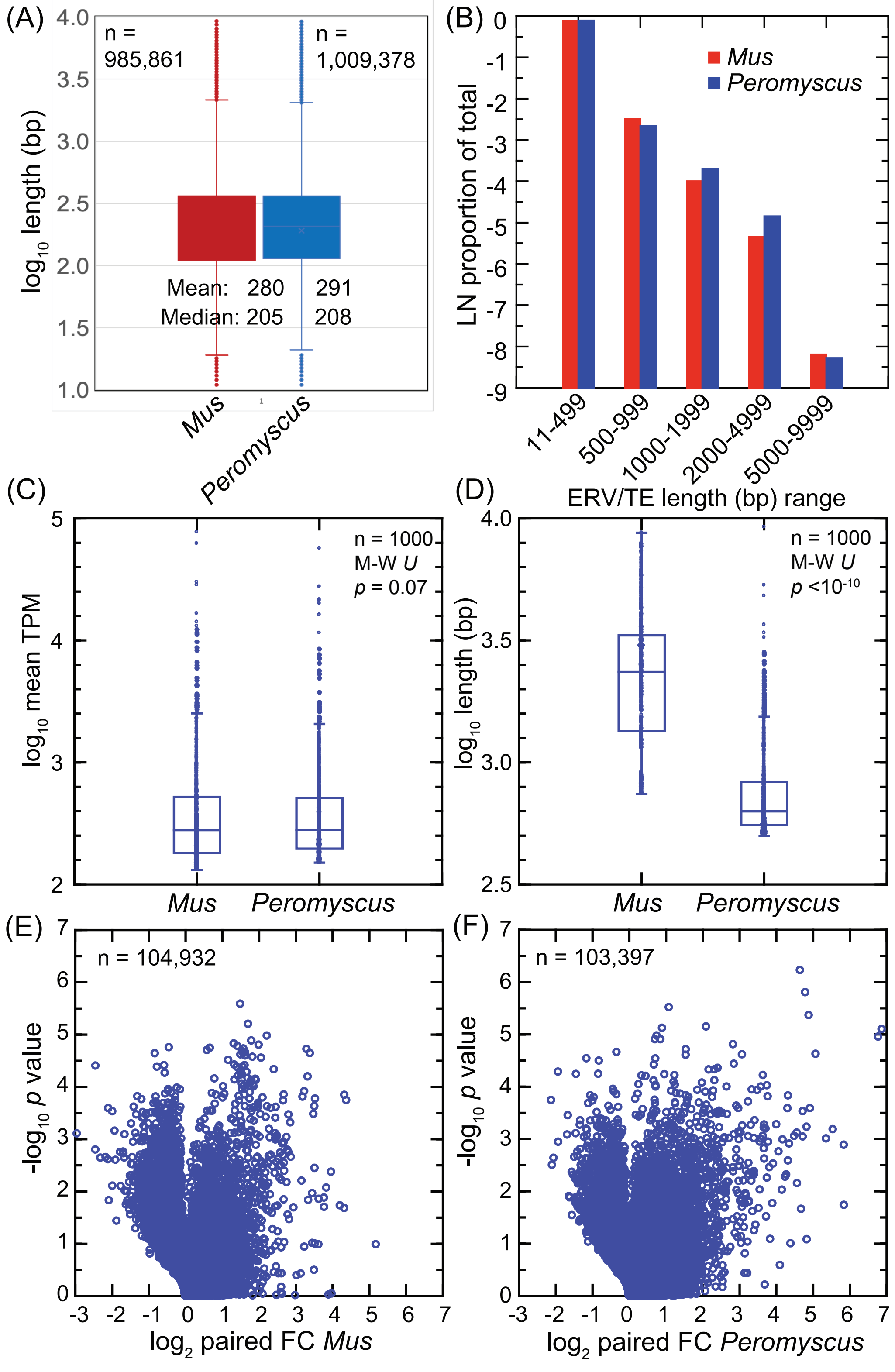


Figure S6. Comparison of ERV/TE sequences of 11 to 9999 base pair (bp) length range in *Mus musculus* and *Peromyscus leucopus* genomes and the transcription of these in low-passage dermal fibroblasts with or without treatment with Pam3CSK4 TLR2 agonist. Panel A is a box-whisker plot of $\log_{10}$-transformed lengths for the entire sets for each species with counts given by "n". Panel B is a graph of natural logarithms (LN) of the proportion of sequences of the specified range (*x*-axis) out of the total number (n) of ERV/TE sequences for each species indicated in panel A. Data for analyses for panels A and B are provided in Dryad Tables D4 (*P. leucopus*) and D5 (*M. musculus*). Panel C ($\log_{10}$ mean TPM values across 5 samples for a sequence for each species) and panel D ($\log_{10}$ length) are box-whisker plots for the top 1000 in descending order of mean TPM values for ERV/TE sequences of 500-9999 bp for *M. musculus* (n=104,932) and *P. leucopus* (n=103,397) for untreated dermal fibroblasts. Data for analyses of panels E and F are from Dryad Table D6. Panels E (*M. musculus*) and F (*P. leucopus*) are volcano plots of $-\log_{10}$ paired *t*-test *p* values (*y*-axis) on $\log_2$-transformed treatment-to-control fold-change (FC) values (*x*-axis) for sequences in range of 500-9999 bp for each species with counts given by "n". Data for analyses of panels E and F are provided in Dryad Tables D8 and D9, respectively.
