## Supplementary material for "Dermal fibroblast cultures recapitulate differences between deermice and mice in responses to a Toll-like receptor agonist": zipped archive of supplemental figures, tables, and text: Supplementary_Text_2.pdf

Descriptions of datasets (sizes) in Excel format spreadsheets at the Dryad (http://datadryad.org) repository (https://doi.org/10.5061/dryad.m905qfvdq):

Dryad Table D1 (5.4 MB). Peromyscus leucopus dermal fibroblast normalized RNA-seq reads per kb of genome-wide CDS by animal source, treatment (control or 1 ¬µg/ml or 10 ¬µg/ml Pam3CSK4), and comparison of 1 ¬µg/ml treatment to control by paired t-test and mean paired fold-change, and mean transcription across all samples by gene

Dryad Table D2 (5.3 MB). Mus musculus dermal fibroblast normalized RNA-seq reads per kb of genome-wide CDS by animal source, treatment (control or 1 ¬µg/ml or 10 ¬µg/ml Pam3CSK4), and comparison of 1 ¬µg/ml treatment to control by paired t-test and mean paired fold-change, and mean transcription across all samples by gene

Dryad Table D3 (2.3 MB). Paired treatment-to-control fold-change (FC) and paired t-tests of transcribed (TPM‚â•10) CDS (n= 14,979) in common for M. musculus (M) and P. leucopus (P) treated with Pam3CSK4 at 1 ¬µg or 10 ¬µg/ml

Dryad Table D4 (46.8 MB). Endogenous retrovirus/transposable element sequences by name, genome location, and length of Peromyscus leucopus LL stock

Dryad Table D5 (38.2 MB). Endogenous retrovirus/transposable element sequences by name, genome location, and length of Mus musculus C57BL/6

Dryad Table D6 (12.4 MB). ERV/TEs ‚â•500 bp of P. leucopus and M. musculus dermal fibroblasts without (control) or with 1 ¬µg/ml Pam3CSK4 and by length, transcription, and fold change of treatment to control

Dryad Table D7 (2.0 MB). Differential gene expression of dermal fibroblasts to TLR agonist by isoform for 13,786 genes of Mus musculus for which there are two isoforms for protein coding sequences

Dryad Table D8 (28.8 MB). Differential experssion of ERV/TEs ‚â•500 bp of Mus musculus (Table D5) in low-passage dermal fibroblasts untreated or treated with 1 ¬µg/ml Pam3CSK4

Dryad Table D9 (35.3 MB). Differential experssion of ERV/TEs ‚â•500 bp of Peromyscus leucopus (Table D4) in low-passage dermal fibroblasts untreated or treated with 1 ¬µg/ml Pam3CSK4
