## Supplementary material for "Dermal fibroblast cultures recapitulate differences between deermice and mice in responses to a Toll-like receptor agonist": zipped archive of supplemental figures, tables, and text: Supplementary_Text_3.pdf

```
python process_fasta.py in.fa out.csv

# process_fasta.py
import csv
import sys
from Bio import SeqIO
from Bio.Seq import Seq
import re

def find_orfs(seq, min_length=30):
    orfs = []
    pattern = 
re.compile(r'([AG][ACGT][ACGT]ATG(?:(?!TAA|TAG|TGA)...)*(?:TAA|TAG|TGA))'
)

    for strand, nuc in [(+1, seq), (-1, seq.reverse_complement())]:
        for match in pattern.finditer(str(nuc)):
            start = match.start()
            end = match.end()
            orf = nuc[start:end]
            if len(orf) >= min_length * 3 and orf[6] == 'G':  # Check if 
the 7th base is G
                if strand == -1:
                    start, end = len(seq) - end, len(seq) - start
                orfs.append((strand, start, end, orf))

    return orfs

def process_fasta(file_path, output_path):
    with open(output_path, 'w', newline='') as out_file:
        csv_writer = csv.writer(out_file)

        # Write header
        csv_writer.writerow(["transcript", "strand", "start", "end", 
"length", "peptide"])

        for record in SeqIO.parse(file_path, "fasta"):
            seq = record.seq
            orfs = find_orfs(seq)

            for strand, start, end, orf in orfs:
                # Adjust start and end positions to exclude the 3 
upstream bases
                adjusted_start = start + 3
                adjusted_end = end

                # Adjust the ORF sequence to exclude the 3 upstream bases
                adjusted_orf = orf[3:]

                peptide = adjusted_orf.translate()
                csv_writer.writerow([
                    record.id,
                    '+' if strand == 1 else '-',
```

```
                    adjusted_start + 1,  # +1 because biological
coordinates are 1-based
                    adjusted_end,
                    len(adjusted_orf),
                    str(peptide)
                ])
if __name__ == "__main__":
    if len(sys.argv) != 3:
        print("Usage: python process_fasta.py <input_fasta>
<output_file>")
        sys.exit(1)

    input_file = sys.argv[1]
    output_file = sys.argv[2]

    process_fasta(input_file, output_file)
    print(f"Results have been written to {output_file}")
```
