## Supplementary material for "Dermal fibroblast cultures recapitulate differences between deermice and mice in responses to a Toll-like receptor agonist": zipped archive of supplemental figures, tables, and text: TableS1.pdf

Table S1. Line listing of samples under BioProject PRJNA1168383 and with their characteristics and accession numbers

| | | | | | Control - saline | | | | Pam3CSK4 1 µg/ml | | | | Pam3CSK4 10 µg/ml | | | |
|---|---|---|---|---|---|---|---|---|---|---|---|---|---|---|---|---|
| number | animal ID | species | stock | sex | BioSample | PE150 reads | SRA | GEO | BioSample | PE150 reads | SRA | GEO | BioSample | PE150 reads | SRA | GEO |
| 1 | 25417 | *Peromyscus leucopus* | LL | female | SAMN45236433 | 62061469 | SRR31668291 | GSM9100090 | SAMN45236432 | 57480172 | SRR31668292 | GSM9100089 | SAMN45236431 | 57802220 | SRR31668293 | GSM9100088 |
| 2 | 25418 | *Peromyscus leucopus* | LL | female | SAMN45236430 | 53885821 | SRR31668294 | GSM9100093 | SAMN45236429 | 62753260 | SRR31668295 | GSM9100092 | SAMN45236428 | 64192593 | SRR31668262 | GSM9100091 |
| 3 | 25459 | *Peromyscus leucopus* | LL | female | SAMN45236424 | 66116931 | SRR31668266 | GSM9100096 | SAMN45236423 | 55069478 | SRR31668267 | GSM9100095 | SAMN45236422 | 71210765 | SRR31668274 | GSM9100094 |
| 4 | 25510 | *Peromyscus leucopus* | LL | male | SAMN45236421 | 68185985 | SRR31668285 | GSM9100099 | SAMN45236420 | 69777800 | SRR31668296 | GSM9100098 | SAMN45236419 | 75123467 | SRR31668297 | GSM9100097 |
| 5 | 25558 | *Peromyscus leucopus* | LL | male | SAMN45236427 | 68264338 | SRR31668263 | GSM9100102 | SAMN45236426 | 67367713 | SRR31668264 | GSM9100101 | SAMN45236425 | 70462671 | SRR31668265 | GSM9100100 |
| 6 | MF1 | *Mus musculus* | CD-1 | female | SAMN45236442 | 70502563 | SRR31668281 | GSM9100069 | SAMN45236441 | 68600982 | SRR31668282 | GSM9100068 | SAMN45236440 | 72470228 | SRR31668283 | GSM9100067 |
| 7 | MF2 | *Mus musculus* | CD-1 | female | SAMN45236445 | 62561601 | SRR31668278 | GSM9100072 | SAMN45236444 | 58171570 | SRR31668279 | GSM9100071 | SAMN45236443 | 59452728 | SRR31668280 | GSM9100070 |
| 8 | MF3 | *Mus musculus* | CD-1 | female | SAMN45236447 | 69456772 | SRR31668276 | GSM9100075 | SAMN45236448 | 62560040 | SRR31668275 | GSM9100074 | SAMN45236446 | 68555317 | SRR31668277 | GSM9100073 |
| 9 | MM1 | *Mus musculus* | CD-1 | male | SAMN45236436 | 63434915 | SRR31668288 | GSM9100078 | SAMN45236435 | 82867487 | SRR31668289 | GSM9100077 | SAMN45236434 | 77940881 | SRR31668290 | GSM9100076 |
| 10 | MM2 | *Mus musculus* | CD-1 | male | SAMN45236439 | 67833497 | SRR31668284 | GSM9100081 | SAMN45236438 | 77880118 | SRR31668286 | GSM9100080 | SAMN45236437 | 60443235 | SRR31668287 | GSM9100079 |
