## Supplementary material for "Dermal fibroblast cultures recapitulate differences between deermice and mice in responses to a Toll-like receptor agonist": zipped archive of supplemental figures, tables, and text: TableS3.pdf

Table S3. Product names of DEGs of CDS among DEGs

| CDS | Product |
|---|---|
| Abcg2 | ATP binding cassette subfamily G member 2 |
| Actb | actin, beta |
| Adamts5 | ADAM metallopeptidase with thrombospondin type 1 motif 5 |
| Adamts9 | ADAM metallopeptidase with thrombospondin type 1 motif 9 |
| Adamtsl2 | ADAMTS-like 2 |
| Ago1 | argonaute RISC catalytic subunit 1 |
| Aqp5 | aquaporin 5 |
| Arg1 | arginase 1 |
| Arg2 | arginase 2 |
| Ass1 | Argininosuccinate synthetase 1 |
| Batf2 | basic leucine zipper transcription factor, ATF-like 2 |
| Bcl3 | B cell leukemia/lymphoma 3 |
| Bdkrb1 | bradykinin receptor, beta 1 |
| Brca1 | breast cancer 1, early onset |
| Bst2 | bone marrow stromal antigen 2 |
| C1r | complement component 1, r subcomponent A |
| C4b | complement C4B |
| Ca3 | carbonic anhydrase 3 |
| Calhm5 | calcium homeostasis modulator family member 5 |
| Casp1 | caspase 1 |
| Cck | cholecystokinin |
| Ccl27a | C-C motif chemokine ligand 27A |
| Ccl7 | C-C motif chemokine ligand 7 |
| Cd248 | CD248 |
| Cd40 | CD40 |
| Cfb | complement factor B |
| Ch25h | cholesterol 25-hydroxylase |
| Chi3l1 | chitinase 3 like 1 (Brp39) |
| Cmpk2 | cytidine/uridine monophosphate kinase 2 |
| Col11a1 | collagen, type XI, alpha 1 |
| Col18a1 | collagen, type XVIII, alpha 1 |
| Col3a1 | collagen, type III, alpha 1 |
| Col4a5 | collagen, type IV, alpha 5 |
| Col5a3 | collagen, type V, alpha 3 |
| Col6a1 | collagen, type VI, alpha 1 |
| Col6a2 | collagen, type VI, alpha 2 |
| Col6a3 | collagen, type VI, alpha 3 |
| Col8a1 | collagen, type VIII, alpha 1 |
| Csf3 | colony stimulating factor 3 (granulocyte) |
| Ctsk | cathepsin K |
| Cxcl1 | C-X-C motif chemokine ligand 1 |
| Cxcl10 | C-X-C motif chemokine ligand 10 |
| Cxcl11 | C-X-C motif chemokine ligand 11 |
| Cxcl12 | C-X-C motif chemokine ligand 12 |
| Cxcl14 | C-X-C motif chemokine ligand 14 |
| Cxcl3 | C-X-C motif chemokine ligand 3 |
| Cxcl5 | C-X-C motif chemokine ligand 5 |
| Dcn | decorin |
| Ddx60 | DExD/H box helicase 60 |
| Dennd3 | DENN domain containing 3 |
| Deptor | DEP domain containing MTOR-interacting protein |
| Dkk2 | dickkopf WNT signaling pathway inhibitor 2 |
| Dpt | dermatopontin |
| Dpy19l2 | Dpy-19 like 2 |
| Dtx3l | deltex 3-like, E3 ubiquitin ligase |
| Ednrb | endothelin receptor type B |
| Efemp1 | EGF-containing fibulin-like extracellular matrix protein 1 |
| Egln3 | Egl-9 family hypoxia-inducible factor 3 |
| Eng | endoglin (CD105) |
| Ereg | epiregulin |
| Fscn1 | fascin |
| Fth1 | Ferritin heavy polypeptide 1 |
| Gbp2b | guanylate binding protein 2b (Gbp1) |
| Gzma | granzyme A |
| H-2_DDa | MHC class I-like protein (XP_028718389.1) |
| H-2_DDa1 | MHC class I-like protein (XP_037055648.1) |
| H-2_LDa-3 | MHC class I-like protein (XP_028718386.2) |
| H-2_Q10a-1 | MHC class I-like protein (XP_037055648.1) |
| H19 | H19, imprinted maternally expressed transcript |
| Hpx | hemopexin |
| Icam1 | intercellular adhesion molecule 1 |
| Icam4 | intercellular adhesion molecule 4 |
| Ifi203 | interferon activated gene 203 |
| Ifi205 | interferon activated gene 205 |
| Ifih1 | interferon induced with helicase C domain 1 (MDA5) |
| Ifngr1 | interferon gamma receptor 1 |
| Igf1 | insulin-like growth factor 1 |
| Ikbke | inhibitor of kappaB kinase epsilon |
| Il11 | interleukin-11 |
| Il15 | interleuki-15 |
| Il1r2 | interleukin-1 receptor, type II |
| Il1rn | interleukin-1 receptor antagonist |
| Il36a | interleukin-36a |
| Il4r | interleukin-4 receptor |
| Il6 | interleukin-6 |
| Il6r | interleukin-6 receptor |
| Insig1 | insulin induced gene 1 |
| Irf7 | interferon regulatory factor 7 |
| Irf9 | interferon regulatory factor 9 |
| Irgm1 | immunity-related GTPase family M member 1 |
| Irgm2 | immunity-related GTPase family M member 2 |
| Ism1 | isthmin 1, angiogenesis inhibitor |
| Kctd12 | potassium channel tetramerisation domain containing 12 |
| Kpna7 | karyopherin subunit alpha 7 |
| Krt14 | keratin 14 |
| Lbp | lipopolysaccharide binding protein |
| Lcn2 | lipocalin 2 |
| Ldoc1 | regulator of NFKB signaling |
| Lpl | lipoprotein lipase |
| Lrig1 | leucine-rich repeats and immunoglobulin-like domains 1 |
| Lrrc17 | leucine rich repeat containing 17 |
| Lum | lumican |
| Ly6e | lymphocyte antigen 6E |
| Mmp12 | matrix metallopeptidase 12 |
| Mmp2 | matrix metallopeptidase 2 |
| Mmp3 | matrix metallopeptidase 3 |
| Mmp9 | matrix metallopeptidase 9 |
| Mt1 | metallothionein 1 |
| Mt2 | metallothionein 2 |
| Mx2 | MX dynamin-like GTPase 2 |
| Myc | myelocytomatosis oncogene |
| Ndn | necdin, MAGE family member |
| Neurl3 | neuralized E3 ubiquitin protein ligase 3 |
| Nfkb1 | nuclear factor of kappa light polypeptide gene enhancer in B cells 1, p105 |
| Nfkbiz | nuclear factor of kappa light polypeptide gene enhancer in B cells inhibitor, zeta |
| Nod1 | nucleotide-binding oligomerization domain containing 1 |
| Nos2 | nitric oxide synthase 2 |
| Oas1 | 2'-5' oligoadenylate synthetase 1 |
| Pglyrp3 | peptidoglycan recognition protein 3 |
| Pla2g2a | phospholipase A2, group IIA |
| Pla2g5 | phospholipase A2, group V |
| Plek | pleckstrin |
| Prrx2 | paired related homeobox 2 |
| Rcan1 | regulator of calcineurin 1 |
| Rgs16 | regulator of G-protein signaling 16 |
| Rgs5 | regulator of G-protein signaling 5 |
| Rigi | RNA sensor RIG-I (Ddx58) |
| Rnf122 | ring finger protein 122 |
| Rsad2 | radical S-adenosyl methionine domain containing 2 (Viperin) |
| Rspo3 | R-spondin-3 |
| Rtp4 | receptor transporter protein 4 |
| S100a8 | S100 calcium binding protein A8 (calgranulin A) |
| S100b | S100 protein, beta polypeptide, neural |
| Saa1 | serum amyloid A 1 |
| Saa3 | serum amyloid A 3 |
| Samd11 | sterile alpha motif domain containing 11 |
| Sdc4 | syndecan 4 |
| Selp | Selectin, platelet |
| Sema3d | Sema domain, immunoglobulin domain (Ig), short basic domain, secreted, 3D |
| Sema7a | Sema domain, immunoglobulin domain (Ig), and GPI membrane anchor, 7A |
| Serpina3f | serine (or cysteine) peptidase inhibitor, clade A, member 3f |
| Serpina3g | serine (or cysteine) peptidase inhibitor, clade A, member 3g |
| Serpina3i | serine (or cysteine) peptidase inhibitor, clade A, member 3I |
| Serpina3n | serine (or cysteine) peptidase inhibitor, clade A, member 3N |
| Slc2a6 | solute carrier family 2, member 6 (GLUT6) |
| Slc7a2 | solute carrier family 7, member 2 (CAT2) |
| Slco2a1 | solute carrier organic anion transporter family, member 2a1 (PGT) |
| Slfn2 | schlafen 2 |
| Slpi | secretory leukocyte peptidase inhibitor |
| Sod2 | superoxide dismutase 2 |
| Sp110 | Sp110 nuclear body protein (Ipr1) |
| Srgn | serglycin (Prg) |
| Stat2 | signal transducer and activator of transcription 2 |
| Stat4 | signal transducer and activator of transcription 4 |
| Sting1 | stimulator of interferon response cGAMP interactor 1 |
| Tagln | transgelin |
| Tfpi2 | tissue factor pathway inhibitor 2 |
| Tgfb1 | transforming growth factor, beta 1 |
| Timeless | timeless circadian clock 1 |
| Tlr2 | Toll-like receptor 2 |
| Tm4sf19 | transmembrane 4 L six family member 19 |
| Tnf | tumor necrosis factor |
| Tnfaip6 | tumor necrosis factor alpha induced protein 6 (TSG-6) |
| Tnfsf11 | tumor necrosis factor (ligand) superfamily, member 11 (RANKL) |
| Tnfsf18 | tumor necrosis factor (ligand) superfamily, member 18 |
| Tnfsf4 | tumor necrosis factor (ligand) superfamily, member 4 |
| Traf1 | TNF receptor-associated factor 1 |
| Trim30d | tripartite motif-containing 30D |
| Tyk | tyrosine kinase 2 |
| Ubd | ubiquitin D (FAT10) |
| Ung | uracil DNA glycosylase |
| Vcam1 | vascular cell adhesion molecule 1 |
| Wnt16 | Wingless-type MMTV integration site family, member 16 |
